# A 10-year longitudinal study reveals bat-driven SARSr-CoV evolution

**DOI:** 10.64898/2026.10.05.754022

**Authors:** Daphné de Riols de Fonclare, Artur Szczepanski, Aleksandra Milewska, Sandrine Corbet, Lucile Rouilly, Laure Diancourt, Baptiste Elie, Estelle Leperchois, Sebastian Lequime, Bryce Leterrier, Sébastien J. Puechmaille, Frédéric Touzalin, Etienne Decroly, Astrid Vabret, Sophie Le Poder, Krzysztof Pyrc, Meriadeg Ar Gouilh

## Abstract

Bats are recognized reservoirs for several zoonoses and have an exceptional tolerance to viruses, likely linked with strong evolutionary selection in immune genes^1^. Notably, in light of the emergence history of severe acute respiratory syndrome coronavirus (SARS-CoV), SARS-CoV-2 is hypothesized to have originated from *Rhinolophus* bats as well^2–6^. In-depth studies are still needed, however, to unravel how SARS-related coronaviruses (SARSr-CoVs) persist and thrive in nature through evolution, virus-host interactions, and cross-species transmission. Here we present a pioneering 10-year longitudinal study of BBCoV-EPI1, a SARSr-CoV in *Rhinolophus* bats that revealed the complexity of virus-host interactions and the tempo at which new viral lineages emerge. Our results demonstrated the impacts of host life cycles — hibernation, female gathering, weaning juveniles, and mating periods — on strong seasonal oscillations in viral prevalence. These pronounced fluctuations drove viral diversification through bottlenecks, founder effects and Darwinian selection, counterbalanced by high mutation rates, fast gradualism, and recombination. We also found that host-cell enzymes are major contributors to BBCoV-EPI1 evolution in bats. The antiviral effect of APOBEC3 detected here in bats is weaker than it is in humans^7^. Conversely, the ADAR enzyme had an unexpectedly strong and beneficial influence on viral evolution, even though the high mutation rate of RNA viruses is usually attributed to viral polymerases’ low replication fidelity^8^. We discovered that key BBCoV-EPI1 innovations in *Rhinolophus* bats were mostly located in proteins involved in cell-cycle hijack and in blocking bat cells’ antiviral response, rather than in the Spike (S)^9,10^. We also showed that the BBCoV-EPI1 S protein efficiently uses angiotensin-converting enzyme 2 (ACE2) receptor to enter *Rhinolophus* bat cells, but not to enter human or sympatric species cells, due to amino-acid differences at the S-ACE2 interface. Last, this work identifies 101 critical amino-acid changes frequently associated with viral fitness shift, and paves the way for future experimental studies focusing on the capacity of SARSr-CoVs to adapt to hosts and cross the species barrier.

## Main

Increased human interference in natural ecosystems has caused a rise in zoonotic spillover and emerging infectious diseases, particularly of RNA viruses like the SARS-related coronaviruses (SARSr-CoVs) that belong to the *Betacoronavirus* genus, and the *Sarbecovirus* subgenus (Fig. 1c)^11^. In the aftermath of SARS-CoV and SARS-CoV-2 emergences, the detection of sarbecoviruses in more than 20 *Rhinolophus* bat species across Asia, Europe, and Africa highlighted their high diversity, adaptability, and capacity to cross species barriers^12^. However, the ecological and molecular processes shaping sarbecovirus evolution in nature remain poorly characterized and understood. Cell entry of sarbecoviruses is primarily mediated by the interaction between the Spike (S) glycoprotein and the host receptor, such as the angiotensin-converting enzyme 2 (ACE2)^13^ for clade 1 (SARS-CoV/SARS-CoV-2) and clade 3, though other factors or co-factors are also involved (*e.g.* TMPRSS2, neuropilin-1, furin, cathepsins, heparan sulfates, etc.)^14^. Moreover, although early 3D structure prediction studies suggested that many mammal species could be infected by SARS-CoV-2^15^, it has been shown that ACE2 sequence similarity alone remains a poor predictor of host susceptibility^16^. Indeed, recent reports showed that viral tropism was rather determined by a limited set of key residues within the receptor-binding domain (RBD) and that viral circulation was strongly restricted by ecological barriers^16,17^. Spillover events do not necessarily result in host jumps, as viral entry must be followed by efficient interspecies transmission to establish new lineages^18^. Understanding how viral genetic diversity arises and expands in nature is therefore critical for assessing emergence risk. While polymerase-induced errors are often invoked to explain viral diversity, the editing activity of deaminase enzymes (APOBEC3, ADAR) and the presence of reactive oxygen species (ROS) can also induce specific mutations on viral genomes, including coronaviruses^19–23^. Such signatures were reported in SARS-CoV-2 genomes^24,25^. However, distinguishing these signals from polymerase-derived changes is challenging. These mutations are heterogeneously filtered by natural selection, depending on protein function and genomic regions. The direction of the selection is estimated by measuring the dN/dS, which is the *ratio* of nonsynonymous (N) to synonymous (S) substitutions^26,27^. The typical observations for coronaviruses combine strong purifying selection (dN/dS < 1) on ORF1a, near-neutral evolution (dN/dS = 1) on ORF1b, and relatively relaxed or diversifying evolution (dN/dS > 1) on structural and accessory genes^28–30^. However, selective pressures at domain- and amino-acid levels remain poorly characterized in coronaviruses in nature. A better understanding of these processes requires a sustained monitoring of viruses within their natural hosts, which remains rare due to the logistical challenges of long-term wildlife studies. Most studies thus rely on short-term or cross-sectional datasets, limiting our understanding of ecological events and molecular mechanisms that shape coronavirus evolution in nature^31,32^. Currently, major knowledge gaps remain regarding the evolution of coronaviruses in nature and the early stages of their emergence, including the dynamics and contribution of recombination, molecular level host-virus interactions, as well as frequency, distribution and fitness impact of mutations. Here we present the results of a decade-long surveillance of BBCoV-EPI1, a SARSr-CoV species circulating in *Rhinolophus ferrumequinum* (great horseshoe bat) in France. The established role of horseshoe bats as key hosts of SARSr-CoVs justified the choice of this species as a natural model to study the ecology and evolution of these viruses in nature. To date, BBCoV-EPI1 is the only *Sarbecovirus* detected in the colony^33^, which is located at the western edge of *R. ferrumequinum’s* distribution range. In order to shed light on the evolution of this bat *Sarbecovirus* and reveal how ecological and molecular processes renew viral diversity in nature, we quantified the effect of virus-host natural interactions and developed a holistic approach combining fieldwork, high-frequency sampling and tailored molecular methods for viral genome detection and sequencing. Using 1,227 BBCoV-EPI1 positive feces collected over ten years, we sequenced 65 BBCoV-EPI1 genomes, each obtained from a unique sample. Then, we studied infection dynamics within the maternity colony, lineages emergence time, selective pressures and evolution rates, and identified 101 nonsynonymous mutations, mostly associated with host deaminase activity, half of which were associated with positive viral fitness. In addition, S-ACE2 interactions were investigated using 3D structural modeling and experimental approaches. Despite the rapid evolution observed in BBCoV-EPI1 and its ability to use *R. ferrumequinum* ACE2 for entry, we found this virus had a low capacity to interact with human ACE2 (hACE2) so far. It suggests limited spillover and emergence risks, contrasting with hACE2 binding and entry capacities of several clade 1 sarbecoviruses found in Asia^6,34^.

**Figure 1.**
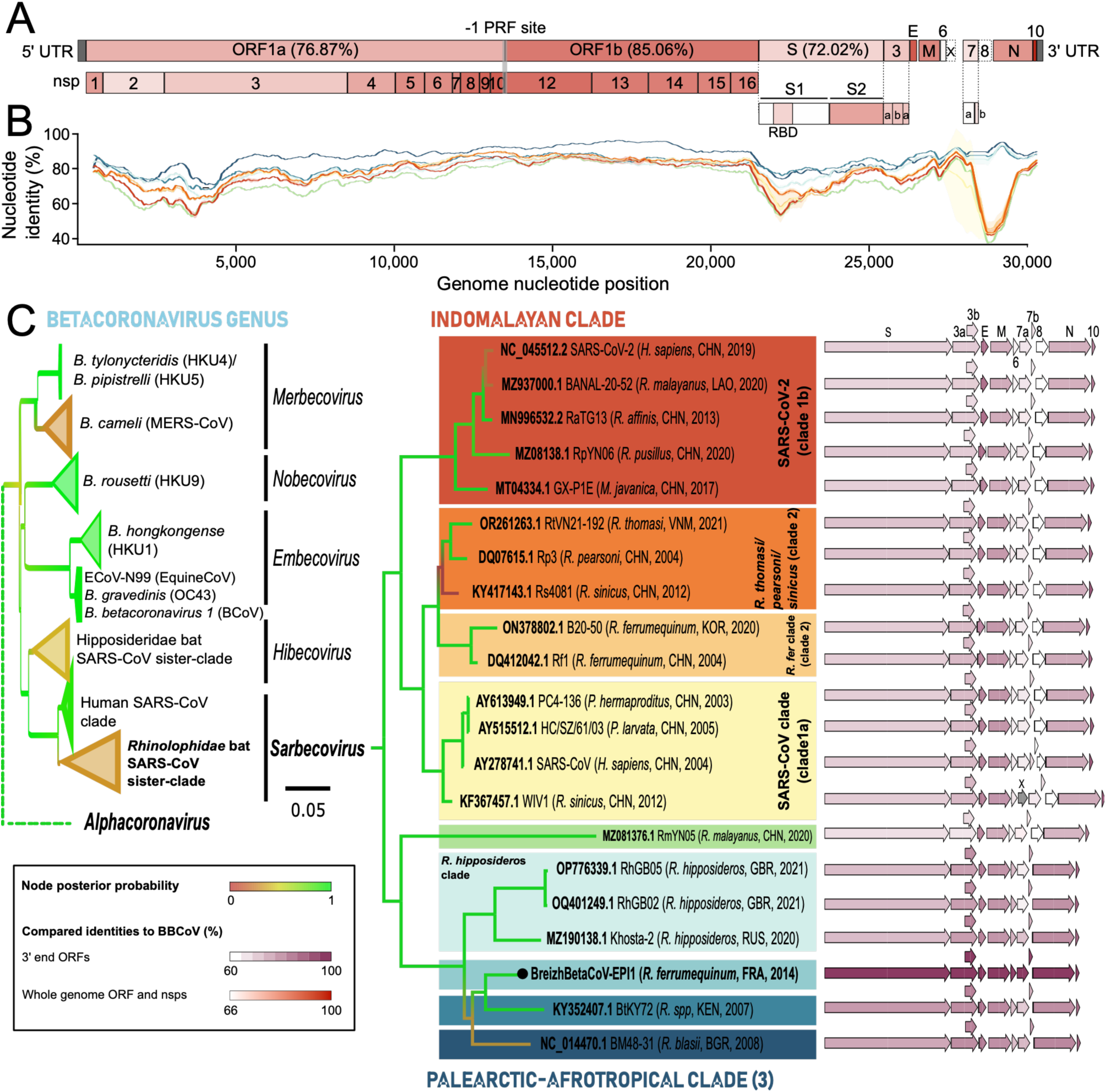
**Compared genomic variability of 21 animal and human SARSr-CoVs sequences**. **A.** The 5’-terminal two-thirds of the genome consists in two large ORFs, 1a and 1b. The presence of a -1 programmed ribosomal frameshift (PRF) site allows the production of the pp1ab polyprotein, further cleaved into 16 non-structural proteins (nsp) involved in viral-host interactions, polyprotein cleavage, genome replication and transcription. The 3’-terminal region comprises genes encoding structural proteins, such as the Spike (S), envelope (E), membrane and nucleocapsid (N) genes. In addition to the genes encoding structural proteins, the 3’-terminal region encompasses accessory genes that are dispensable for virus replication. The ORFs and nsps are colored following the MPNSI (%) of the considered sarbecoviruses relative to BBCoV. **B.** Identity plot of predefined *Sarbecovirus* clades, colored as in Fig. 1c, and using BBCoV-EPI1-P10-G05 (2014) as the reference. **C. Left:** RNA-dependent RNA-polymerase (RdRp)-based phylogeny of the *Betacoronavirus* genus highlighting sub-genus relationships, with collapsed clades and branches colored following the nodes’ robustness (posterior probability). **Right:** Phylogenetic reconstruction of the 3’-end of 21 SARSr-CoVs genomes. Indomalayan clades are shaded in red, palearctic-afrotropical clades are shaded in blue, and ambiguous clade RmYN05 is shown in green. The 3’-end of the *Sarbecovirus* genomes is also represented, with each gene colored according to its percentage identity to BBCoV-EPI1.

## BBCoV-EPI1 is a clade 3 SARSr-CoV

Among 65 BBCoV-EPI1 genomes sequenced over ten years, we used the 29,264-nt P10-G05 (2014) as the reference. Its closest known relative was BM48-31 (86.26% nucleotide identity), sampled in Bulgaria (Figs. 1a, 1b, Supplementary Fig. 1 and Supplementary Table 1)^35^. We then compared P10-G05 with a representative subset of 20 animal and human *Sarbecovirus* genomes from the NCBI Virus database. Phylogenetic analyses of the 3′-end region places BBCoV-EPI1 within the Palearctic-Afrotropical clade, known as clade 3 (Fig. 1c). Its receptor-binding motif (RBM) is most closely related to those of clade 3 sarbecoviruses Khosta-2 (Russia, 2020) and BtKY72 (Kenya, 2007) (Supplementary Fig. 2)^36,37^. Intriguingly, highly divergent RmYN05 (China, 2020), occupies an intermediate position between the Indomalayan (clades 1 and 2) and the Palearctic- Afrotropical (clade 3) clades, rooting the latter (Fig. 1c)^38^. This branching may either result from a S gene affected by a shared ancestry with long-distance lineage diffusion, convergent evolution, or an ancient recombination event, each scenario offering critical insights into the deep origin of both the Palearctic-Afrotropical clade and BBCoV-EPI1. Supporting the first scenario, where *Sarbecovirus* diversity originated in Asia and spread westward from the common ancestor it shares with RmYN05, are the overall conserved 3’end gene synteny and the loss of the ORF8 in all members of the Palearctic-Afrotropical clade following their divergence from RmYN05 (Fig. 1c). These repeated losses, driven by different mechanisms, support a scenario of convergent evolution targeting this gene. While ORF8 deletion in SARS-CoV-2 importantly correlates with attenuated symptoms^39–41^, its functional impact in animal sarbecoviruses remains unresolved^42^. Importantly, shared ancestry and convergent evolution are not mutually exclusive; their interplay over time likely contributed to the complex evolutionary origin of the Palearctic-Afrotropical clade and of BBCoV-EPI1.

## Variations in circulation are driven by life history traits of bats

Beyond its phylogenetic origin, understanding the circulation dynamics of BBCoV-EPI1 is critical for assessing its potential impact and elucidating its evolutionary trajectory. To this end, we conducted a longitudinal survey, collecting 2,885 fecal samples from a *R. ferrumequinum* maternity colony in Brittany (western France), over a decade (2014 to 2024). Samples were screened using a RT-qPCR system specific to the *Sarbecovirus* envelope gene^43^, enabling an estimation of viral prevalence at each time point. Over the study period, 42.5% (1,227/2,885) of the samples were found positive for BBCoV-EPI1 (Fig. 2a and Supplementary Tables 2 and 3), as confirmed by sequencing of PCR products and viral genomes. Prevalence and excretion respectively spanned from 0 to 97.9% (Fig. 2a), and from 1.31 to 3.7x10^9^ copies/µL (Fig. 2b). According to these variations, each year was divided into “Low” (May-July), “Intermediate” (August) and “High” (September-April) prevalence periods (Figs. 2a, 2b and Supplementary Table 3). Notably, a rise in excretion levels precedes increased prevalence by one month, and the two are strongly correlated (Spearman, r = 0.70; p = 1.56 × 10⁻⁵). This supports the hypothesis that once the epidemic begins, early infections with elevated shedding dominate detected cases, thereby increasing the likelihood of detecting higher excretion level in addition to further amplifying transmission, as observed for SARS-CoV-2^44^. We postulate that these variations in prevalence and excretion are driven by bats’ life history traits, particularly the alternation between seasonal movement, torpor (winter) and weaning (June-July). To further investigate these dynamics within the colony, we formulated two mathematical transmission models using Monolix software: a susceptible-infected-recovered-susceptible (SIRS) model, and a MSIRS model, that includes a M compartment representing naive and maternal-antibody-protected juveniles (Supplementary Methods 1). The best-fitting MSIRS model yielded daily transmission (β = 0.37), recovery (γ = 0.005), immunity loss (δ = 0.021), and maternal antibody loss (σ = 0.1). We found that model fitting improved when adding the M compartment, which shows that the primary infections of juveniles after weaning in early July and the associated waning of maternally-induced immunity both likely contribute to increased excretion and subsequent epidemic surge in August (Supplementary Table 4).

**Figure 2.**
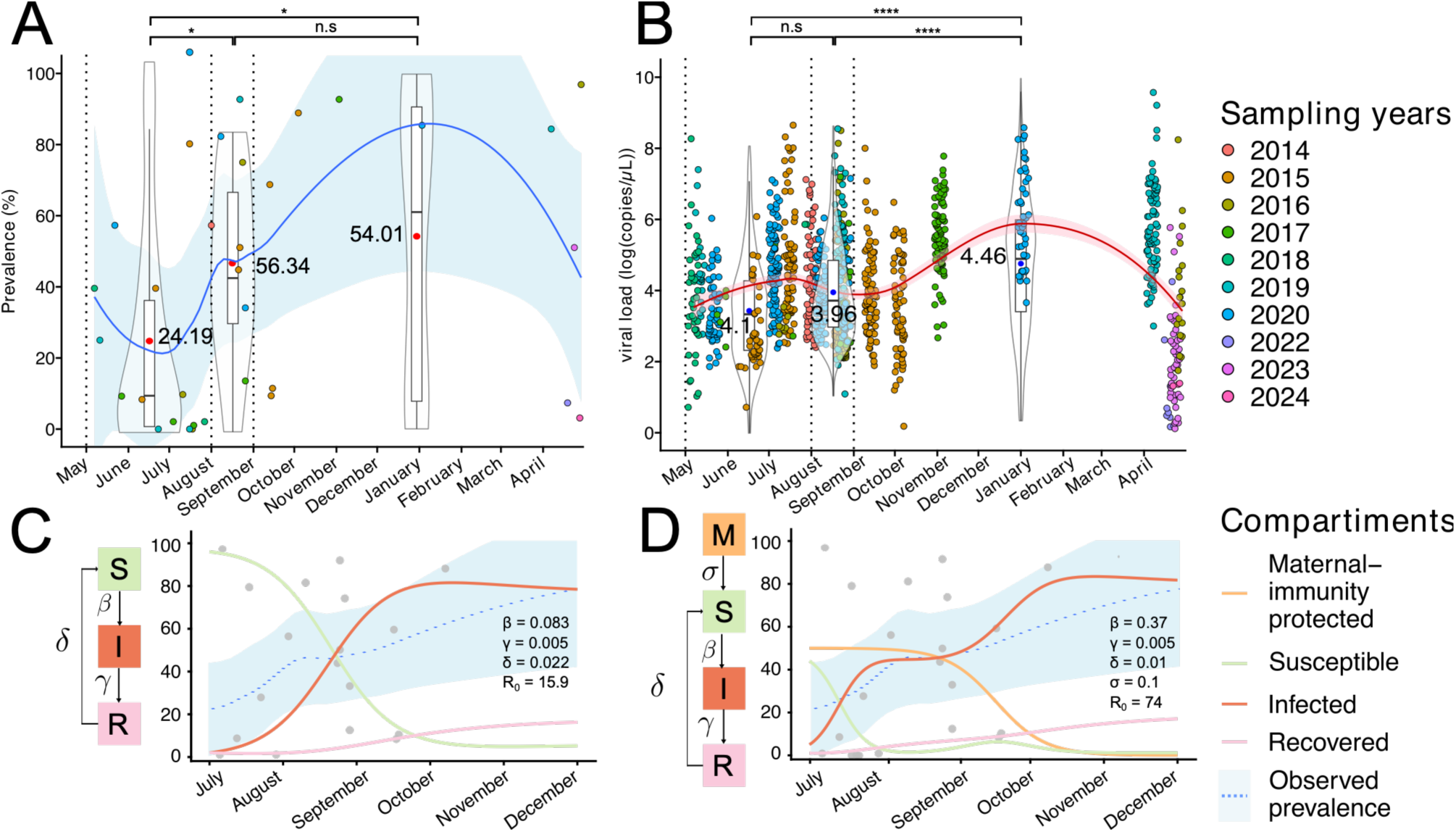
A, B. Annual prevalence and excretion dynamics of BBCoV-EPI1 circulation in the *R. ferrumequinum* colony. Dotted lines divide time into biological seasons ("Low", "High", and "Intermediate" viral prevalence) **A.** One point stands for the prevalence estimated from a single 96-well plate of samples. A LOESS regression curve is shown in blue, with the 95% confidence interval shaded in light blue. The mean prevalence for each biological season is indicated by a red dot inside the boxplot, surrounded by a violin plot indicating the distribution of values. **B.** One point stands for one measure of individual excretion. A LOESS regression curve is shown in red, with the 95% confidence interval shaded in light red. The mean excretion for each biological season is indicated by a blue dot inside the boxplot, surrounded by a violin plot indicating the distribution of values. **C,D. SIR (C) and MSIRS (D) models of BBCoV-EPI1 epidemiology dynamics in the colony.** Susceptible individuals are shown in green, infected in red, recovered in pink, and maternal-immunity protected juveniles in orange.

## Drivers of BBCoV-EPI1 evolution

We obtained 65 full-length BBCoV-EPI1 genomes through deep-sequencing, which we used to infer Bayesian phylogenies for both the whole genome and non-recombinant regions (Figs. 3a, 3d). Over the decade-long study period, BBCoV-EPI1 evolved at a rate of 1.686 × 10⁻³ substitutions/site/year (s/s/y), one of the highest evolutionary rates ever observed for coronaviruses. This rate contrasts with SARS-CoV-2’s inter-lineage rate of 7 × 10⁻⁴ s/s/y^42^. Notably, divergence rates were heterogeneous across genes and over time, with a marked increase observed from 2020 onward (Fig. 3b). From 2014 to 2019, new variants emerged gradually, followed by a genetic sweep in 2020 marked by the rise of a super-clade derived from 2015 lineages. This pattern suggests circulation outside the colony between 2016 and 2020, with subsequent reintroduction (Fig 3a). In light of these results and given that coronaviruses frequently recombine^45^, resulting in a mosaic genome structure with distinct evolutionary history for individual genes, we screened for recombination events and identified 8 putative breakpoints associated with four major events (Fig. 3c and Supplementary Table 5). A notable example is 2020B clade, which inherited its distal genomic region (including the S gene) from progeny of a 2016 lineage. Recombination requires coinfections in individual bats, implying the simultaneous circulation of imported lineages within the colony. *R. ferrumequinum* females are philopatric^46^, but their movements between sites, potentially driven by reproductive behavior or predator-induced disturbances, may have facilitated viral diffusion. These factors likely interleaved to promote spatial dispersion of bats in 2015, enabling the spread of viral lineages outside the colony. Under this hypothesis, after five years of evolution, the reintroduction of progeny viruses in 2019-2020 led to co-infections with local lineages and recombination. This is further supported by evidence of bat disturbances—owl (*Tyto alba*) pellets and beech marten (*Martes foina*) feces—found at the study site in 2015. These observations align with recalculated time to most recent common ancestor estimates of 2020 and 2016 lineages, which also point to a lineage split in 2015 (Fig. 3a). These findings enhance our understanding of the drivers, the frequency and location of recombination in a specific *Sarbecovirus* lineage over ten years in nature. Non-recombinant phylogenies displayed variable divergence rates, reflecting different evolutionary constraints among genes (Fig. 3d). Lowest divergence rates were found in early ORF1a and ORF1b (regions 1 and 3-5), aligning with strong functional constraints related to essential roles in the viral cycle (Fig. 3d). Moderate rates were observed at the 3′ end of the genome (regions 6 & 8, S1-RBD), reflecting a balance between structural constraints and surface exposure of these proteins, potentially driven by immune-escape pressures. Unexpectedly, the highest rates were detected in late ORF1a (region 2, nsp2-8), suggesting either relaxed structural constraints or distinct selective pressures acting on these nonstructural proteins (Fig. 3d).

**Figure 3.**
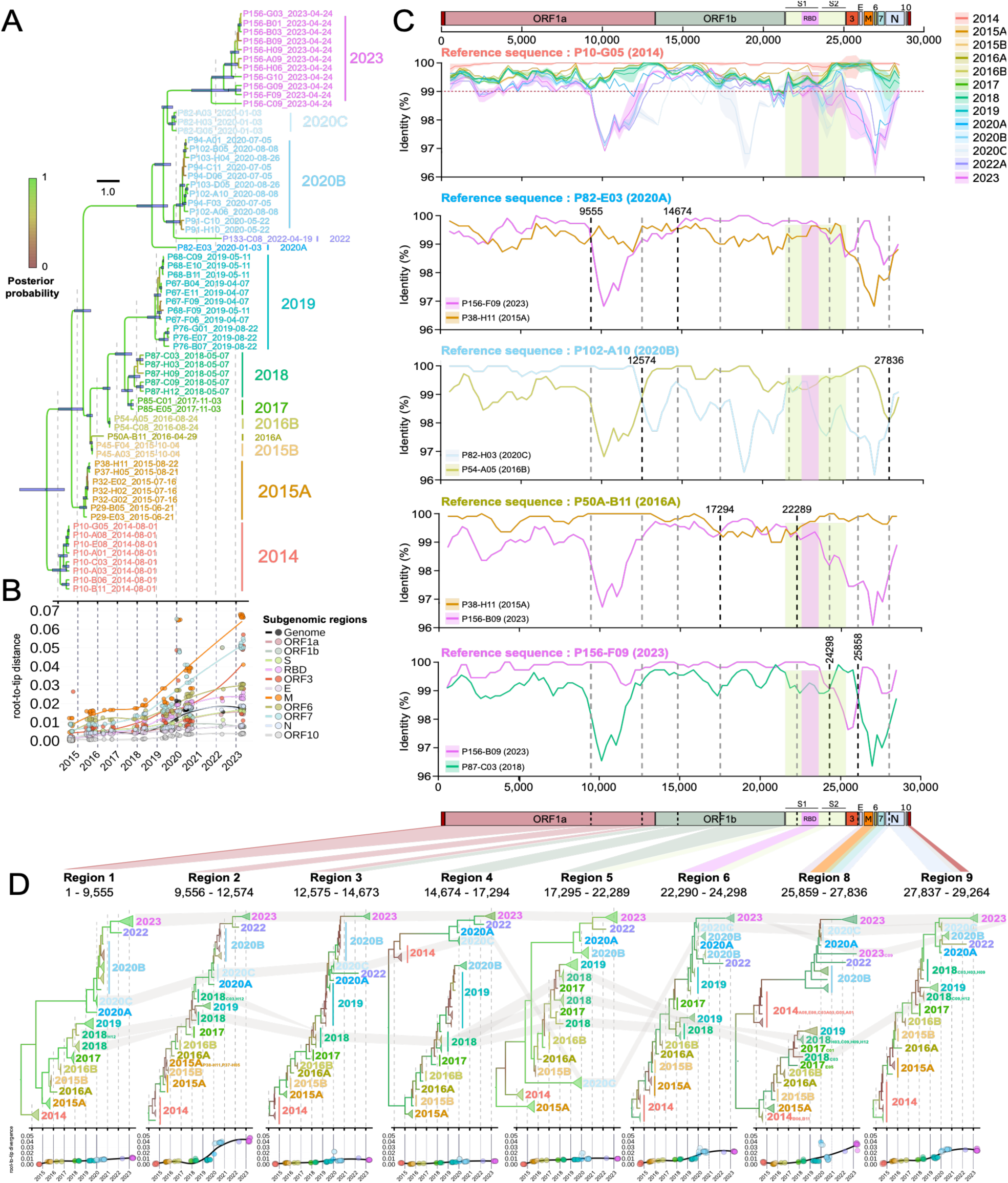
Phylogenetic analyses of 65 BBCoV-EPI1 genomes. **A**. Bayesian phylogeny of 65 BBCoV-EPI1 genomes sampled between 2014 and 2024. Node support (posterior probability) is shown as a color gradient along the branches. **B.** Root-to-tip analyses showing the progressive accumulation of substitutions per site (s/s) since 2014, estimated for full genomes and for subgenomic regions. Each point represents one region. **C**. Identification of four major recombination events using RDP4. Identities of the consensus sequences of each clade relative to BBCoV-EPI1-P10-G05, and of the recombinant sequence (used as the reference) relative to its parental genomes for each recombination event, are shown. **D.** Phylogeny of non-recombinant regions. The phylogeny of region 7 is not shown due to convergence issues and low node support (Supplementary Fig. 3). Root-to-tip distances for each region are shown at the bottom of each tree. Each point, colored by clade, represents the divergence value of a region for each genome.

To further characterize nucleotide diversity and variation trends, we computed substitution rates of each genome and sub-genome (Supplementary Table 6). Here we suggest that the above-average increase of diversity observed in 2015 and 2020, in genomes and several sub-regions (Fig. 4a), may be related to host movements, and to recombination following reintroduction, respectively. To determine the nature of the selection, we computed dN/dS *ratio* at several scales. Sliding window analyses showed that, on average, ORF1a and ORF1b were primarily subjected to purifying and neutral selection, respectively (Fig. 4b). This aligns with their critical roles in viral-host interaction and genome replication, which require structural and functional conservation. Contrastingly, the 3′-end of the genome was mainly targeted by diversifying selection (Fig. 4b), as expected for genes encoding surface proteins exposed to host immunity. Regarding temporal trends, Nsp2, nsp3/PL^pro^ and nsp4 of ORF1a exhibited transient diversifying selection in 2015, 2018, and 2023, respectively (Fig. 4c and Supplementary Table 7). Given the role of these proteins in cell cycle, transient positive selection may reflect an active adaptation towards enhanced replication and host immunity shut-down. Additionally, the substitution rate of these nsps is lower than average, and the positive selection affecting them likely mirrors a bottleneck effect acting on low genetic diversity. Alternatively, strong positive selection, occurring against a backdrop of purifying selection (*i.e* in a highly constrained fitness landscape) could accelerate the selection of beneficial variants capable of counteracting innate immunity, promoting viral replication and facilitating spread into new lineages (Fig. 4d (up)). Also, the detection of numerous positively selected proteins associated with minority variants suggests that only a fraction of these mutations are ultimately selected as major variants (Fig. 4d (down)). This highlights the potential impact of minority variants on viral fitness and underscores their role as a reservoir for adaptive evolution. We observed that majority consensus and intra-host diversity (minor variants) variations trends were opposed to and aligned with those of prevalence, respectively (Fig. 4e). Moreover, nsp2, nsp3/PL^pro^ and nsp4 were submitted to positive selection pressure between late April and mid-July, when females are clearing the virus and its circulation is either declining or very low (Fig. 4d, 4e and Supplementary Table 7). This illustrates a clear seasonal trend of this strong selective pressure on the virus preceding or coinciding with a bottleneck effect occurring in May-July. Then, low prevalence in early summer leads to a rise in relative diversity, from which better-fitting variants may largely contribute to the epidemic surge in July-August by a founder effect^47^ (Fig. 4e (up)). In turn, the primary infection of naive juveniles (accounting for 50% of the maternity colony in August) could fuel the strong increase in viral prevalence and excretion, in mid-summer, that lasts for several months and promotes females’ reinfection with new variants in autumn, before they move to mating sites.

**Figure 4.**
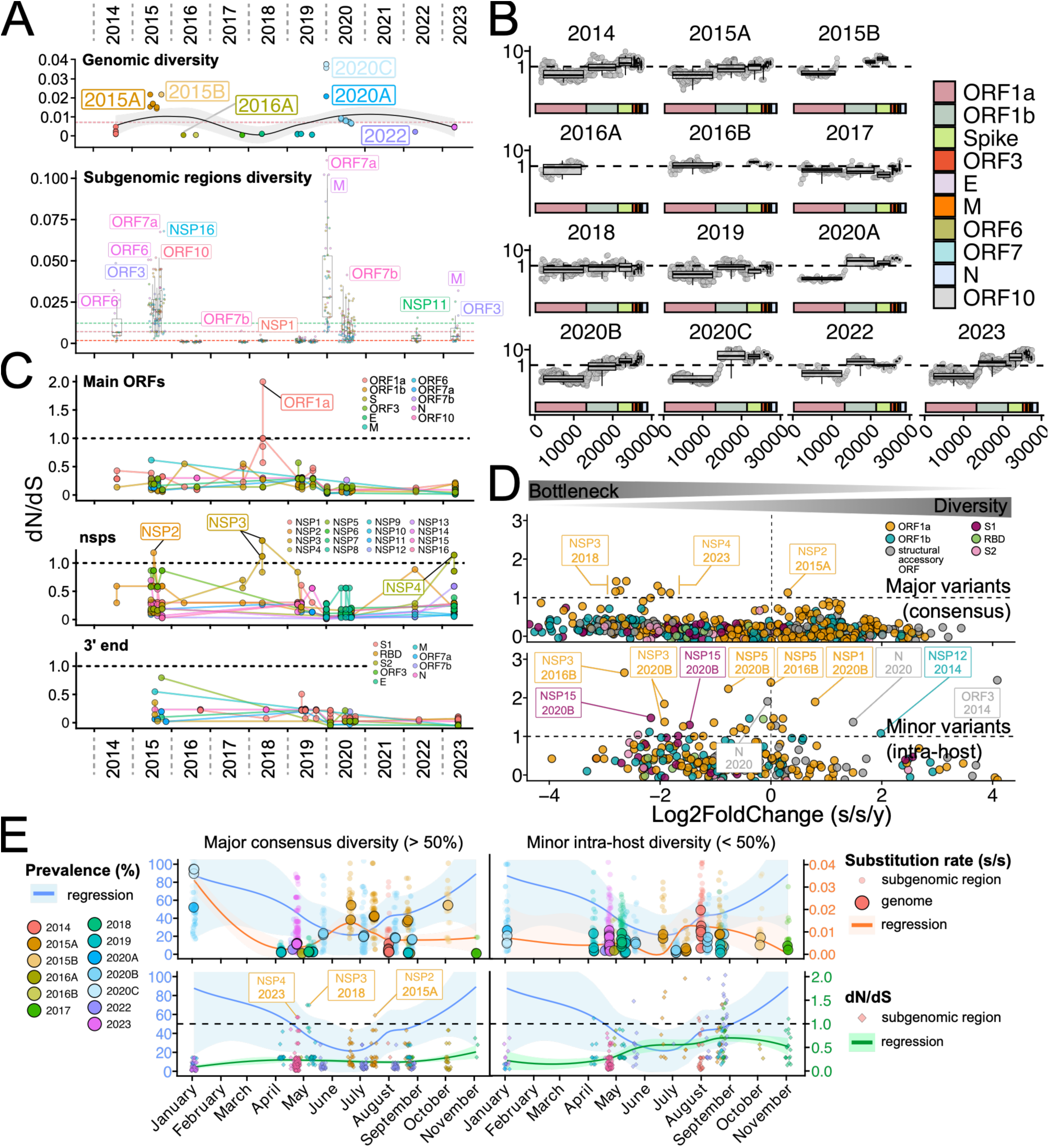
Diversity, selective pressure, and prevalence. **A.** Genomic (up) and subgenomic (down) substitution rates by clade, across time. Pink dashed line indicates the mean genomic substitution rate (7.13 x 10^-3^ s/s/y), green dashed line the mean subgenomic substitution rate (1.22 x 10^-2^ s/s/y), and the red dashed line the BEAST-estimated (GTRIG model) mean genomic substitution rate (1.69 x 10^-3^ s/s/y). Boxplots indicate mean substitution rates per subgenomic regions. **B.** Sliding-window of selective pressure (dN/dS) by clade and over the genome (window = 3000, step = 300). Boxplots indicate mean substitution rates per subgenomic regions **C.** Temporal changes in dN/dS across ORF, nsps (top) and structural/accessory proteins (down) **D.** DN/dS calculated on subgenomic regions and plotted as function of substitution rate (s/s/y), expressed as log2 fold-change, for major consensus (up) and minor intra-host (down) variants. Vertical and horizontal dashed lines represent the mean substitution rate threshold for each nsp and the dN/dS = 1 threshold, respectively. **E.** Annual substitution rates (up, circles) and selective pressure dynamics (down, losanges), for major consensus (left) and diversity (right), both colored according to their clade.

## Host enzymes shape the mutational landscape

We examined the mutational *spectrum* of BBCoV-EPI1 and found that mutations compatible with ADAR (A → G, T → C) and APOBEC3 (C → T) activities accounted for 77.1% of all mutations (Fig. 5a). This suggests that host editing enzymes play a major role in shaping the genetic diversity of BBCoV-EPI1 genomes in bats. To minimize the possible confounding effects of selection, we further explore the nucleotide variability by analyzing the mutational *spectrum* at 4-fold degenerate sites, where any possible changes are synonymous. We then compared this spectrum to that of SARS-CoV-2 in humans, and found that C → T, G → T, and G → A mutations predominated in both host-virus contexts (Fig. 5b and Supplementary Tables 8-11). In bats, BBCoV-EPI1 showed a 59% increase in the rate of ADAR-associated mutations, and a 35% decrease of APOBEC-associated mutations, when compared to SARS-CoV-2 in humans (see Supplementary Fig. 4 for representation of the fraction of all mutations). This suggests that, in *R. ferrumequinum*, ADAR exerts a stronger mutational effect while effect of APOBEC3 is weaker. Moreover, *R. ferrumequinum* showed fewer ROS-linked G → T mutations than humans, a pattern consistent with bats’ evolutionary adaptations to flight, which likely mitigate the oxidative stress associated with their flight-induced high metabolic rate^48,49^. The symmetric proportion of complementary mutation pairs (A → G / G → A; T → C / C → T) likely mirrors a dynamic evolutionary *equilibrium* (Figs. 5a, 5b). This balance may indicate a functional trade-off between the effects of APOBEC3, ADAR, and other mutational sources, which collectively preserve genomic stability by maintaining a viable GC content while fostering adaptability through the frequent emergence of genetic innovations. Next, we assessed the fitness effect of nonsynonymous mutations, adapting a pipeline originally conceived for SARS-CoV-2^50^. Over the decade-long study period, 55 of the 101 nonsynonymous mutations identified were mainly ADAR-associated and linked to a positive viral fitness effect (Fig. 5c). The relatively high frequency of such mutations (5 per year, on average), likely facilitates viral adaptation to changing contexts and increases the likelihood that multiple lineages will co-circulate. The primary contributor to enhanced fitness and, thus, to diversifying viral evolution, included 33 mutations in nsp1-10 of ORF1a. This further underscores the pivotal role of these viral proteins in modulating host cellular machinery and counteracting innate immune response (Fig. 4c). We also observed sporadic reversions and non-fixed mutations across several clades (Fig. 5c), most of which were associated with negative fitness effects. In addition, ten nonsynonymous mutations were identified in the S, predominantly within the NTD, with further substitutions in the RBM and near the S1/S2 cleavage site (Figs. 5c, 6a, 6b and Supplementary Tables 12 and 13).

**Figure 5.**
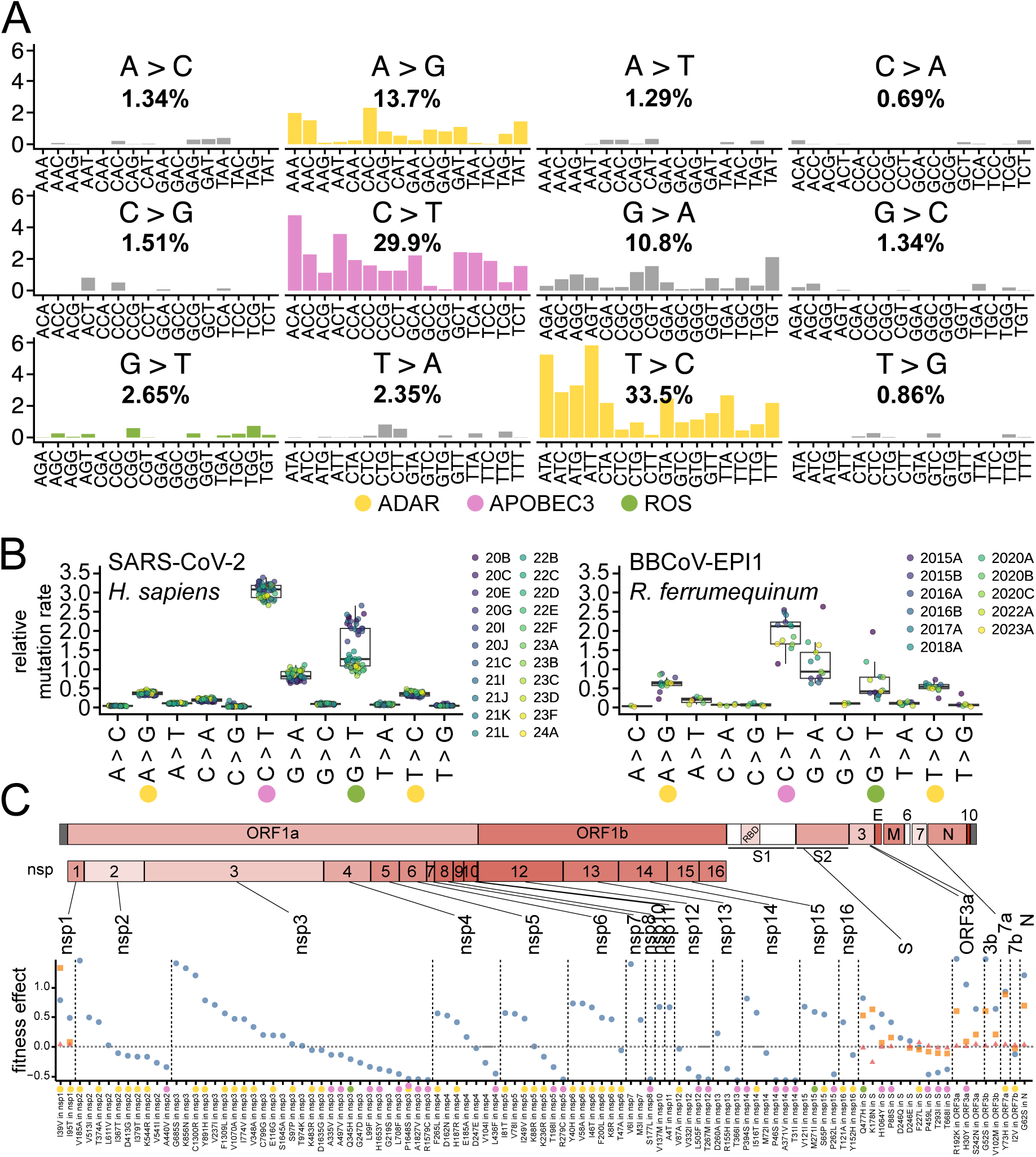
Mutational landscape of BBCoV-EPI1, and estimation of its impact on genomic fitness. **A.** Proportion of the 12 substitution types. For each, its overall proportion in the entire dataset is indicated in bold beneath the substitution symbol. The histograms show the distribution of each mutation type as a function of the flanking nucleotides and are colored according to the associated mutational process: ADAR (yellow), APOBEC3 (pink), or ROS (green). **B.** Relative rates of neutral mutations of BBCoV-EPI1 and SARS-CoV-2 clades, estimated using 4-fold degenerate synonymous sites and normalized to nucleotide composition. **C.** Fitness effects of amino-acid mutations fixed in at least one clade, ordered by decreasing fitness values for each ORF and nsp. The point shapes indicate the fitness estimates, based on: (i) the entire dataset (blue circles); (ii) forward mutations newly acquired in clades that did not previously carry the mutation (orange squares); (iii) reversions in clades where the mutation is fixed (red triangles). Amino-acids changes are represented by circles, colored according to the mutational process: ADAR (yellow), APOBEC3 (pink), or ROS (green).

**Figure 6.**
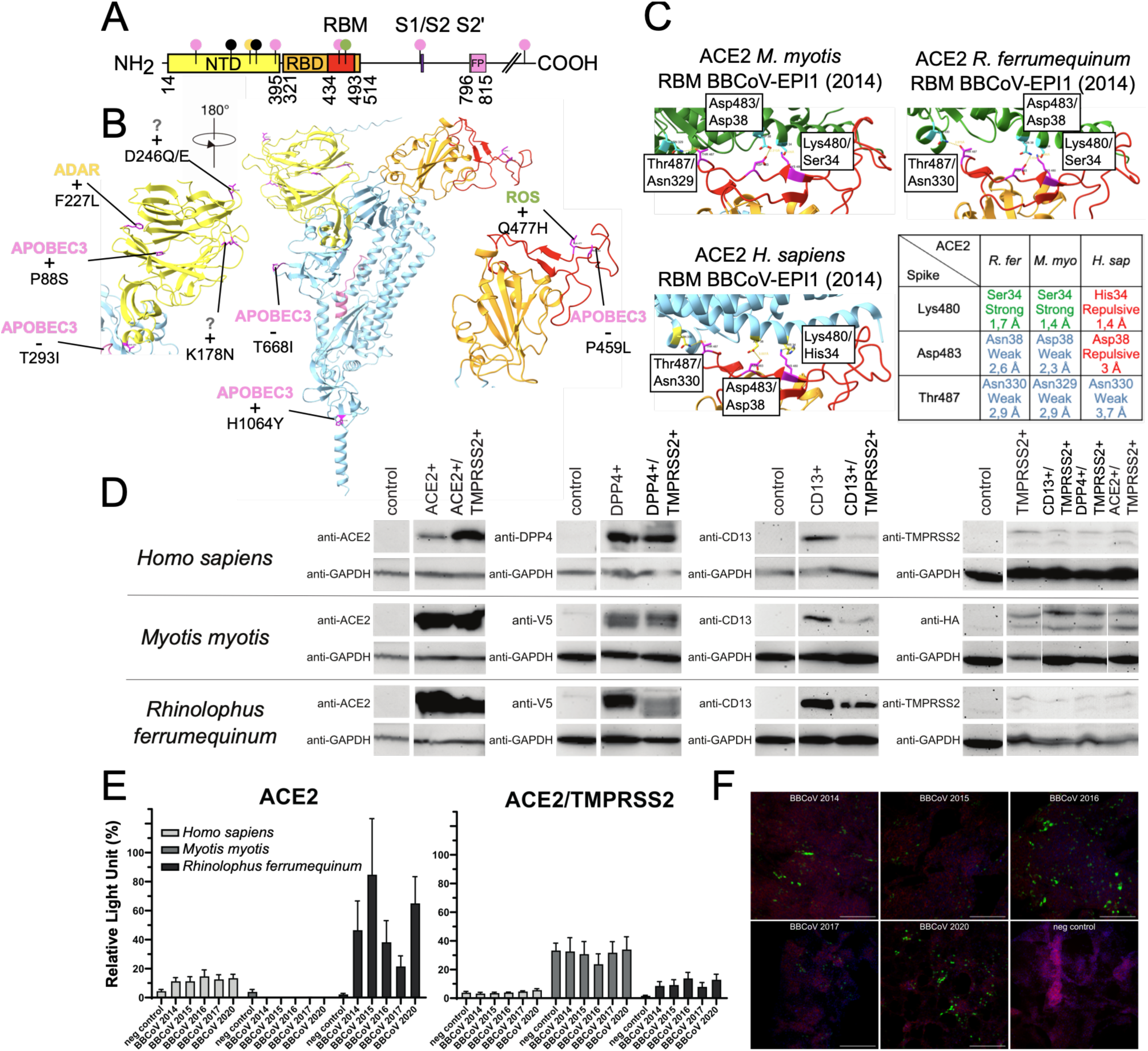
Dissection of BBCoV-EPI1 S-ACE2 affinity using computational modeling and experimental approaches. **A.** BBCoV-EPI1 S protein domain organization showing the NTD (yellow), RBD (orange), RBM (red), S1/S2 cleavage site (violet), S2′ cleavage site (green), and fusion peptide (FP, pink). Mutations detected in this study are indicated and colored according to the associated mutational processes: ADAR (yellow), APOBEC3 (pink), or ROS (green). **B.** Three-dimensional structure of an S protein monomer with enlarged views of the NTD (left) and RBD (right). The RBM (red) is located within the RBD (orange). Detected mutations are mapped on the structure; plus or minus signs indicate positive or negative fitness effects, respectively. The associated antiviral enzyme is depicted. **C.** Identification of S residues from BBCoV-EPI1-P10-G05 potentially involved in ACE2 binding, according to species. **D.** Protein expression analysis in modified A549 cells transduced twice with lentiviral vectors. After antibiotic selection, cell lysates were analyzed by SDS-PAGE and Western blotting. **E.** A549 cells expressing ACE2 alone (left) or ACE2 with TMPRSS2 (right) were infected with HIV pseudoparticles pseudotyped with BBCoV-EPI1 S proteins, bearing different mutations and derived from consensus sequences from 2014, 2015, 2016, 2017, and 2020. **F.** At 72 h post-infection, cells were fixed and stained: the Venus fluorescent protein carried by the pseudoviruses is shown in green, actin in red, and DNA in blue.

## BBCoV-EPI1 uses ACE2 to enter cells

Building on our analysis of S protein evolution (Figs. 5c, 6a, 6b and Supplementary Tables 12 and 13), we investigated the receptor usage, mutation-dependent entry efficiency, and zoonotic potential of BBCoV-EPI1 through a combination of computational modeling and experimental assays. We first modeled the structure of the S protein and its interaction with ACE2 receptors from sympatric bats species and human, using AlphaFold3. Globally, BBCoV-EPI1 S exhibited a similar organization to those of SARS-CoV (RMSD = 2.22Å) and SARS-CoV-2 (RMSD = 4.42Å). Unlike SARS-CoV-2, BBCoV-EPI1 S does not possess a canonical furin recognition motif at the S1/S2 cleavage site and also lacks a known TMPRSS2 recognition site. Instead, it displays monobasic sites at S1/S2 (TNTR_669_) and S2’ regions (HR_796_) (Fig. 6a). Monobasic sites are cleaved by endosomal cathepsins, which are associated with endocytosis-mediated entry, a pathway that favors enteric tropism. This observation supports the hypothesis that BBCoV-EPI1 exhibits enteric tropism. We also found that BBCoV-EPI1 S formed strong ionic interactions with *R. ferrumequinum* ACE2, due to opposite charges and close interactions of key amino-acids (Fig. 6c). These interactions appear permissive in *M. myotis*, but repulsive in *Homo sapiens*. Pseudoviruses bearing BBCoV-EPI1 S proteins, further supported modeling findings. In turn, *R. ferrumequinum* ACE2 supported higher entry efficiency than ACE2 from *M. myotis*, *H*. *sapiens,* and than other putative receptors, such as dipeptidyl-peptidase-4 (DDP4) and cluster of differentiation 13 (CD13), both with or without trypsin (Supplementary Figs. 5 and 6). Regarding the influence of S protein mutations on entry efficiency, pseudoviruses bearing different S proteins exhibited contrasted normalized luminescence. After a significant increase between 2014 and 2015, entry efficiency declined progressively from 2015 to 2017, remained comparably low in 2019, and rebounded markedly in 2020 (Figs. 6e, 6f and Supplementary Tables 14 and 15). These year-to-year fluctuations in entry efficiency aligned with the evolutionary dynamics observed over time. Notably, the 2015 peak in selective pressure coincides with the emergence of the beneficial D246E/Q mutations and the resulting increased entry efficiency compared to 2014 (Figs. 5c, 6a, 6b, 6e). Subsequently, the accumulation of two deleterious mutations (T293I, T668I) during the gradual decline in entry efficiency in 2016 and 2017, contrasts with the contemporaneous emergence of two beneficial mutations (D246E, K178N) (Figs. 6a, 6b and Supplementary Table 12). T293I contributes to the S2 interaction interface and T668I lies upstream of the S1/S2 cleavage site (Figs. 6a, 6b and Supplementary Tables 12 and 13). In 2020, additional beneficial innovations emerged (P88S, Q477H, H1064Y, D246Q), restoring and enhancing entry, despite the persistence of previous deleterious mutations. Intriguingly, the presence of TMPRSS2 significantly reduced entry efficiency (Fig. 6e (right) and Supplementary Fig. 8). Potential explanations include lower ACE2 levels in double-transfected cells, cellular stress induced by protease overexpression, or the effect of protease when released into the medium following autocleavage. Furthermore, the monobasic sequence at the S2′ site likely reduces protease-mediated cleavage. TMPRSS2 co-expression may also interfere with the NTD, which participates in RBD dynamics and potentially interacts with alternative receptors, thereby impairing the ACE2-mediated entry.

## Discussion

The emergence of SARS-CoV, in 2002, triggered a burst of interest in sarbecoviruses present in bat reservoirs. However, the long-term evolution dynamics of these viruses in nature remain largely undescribed. Through a pioneering 10-year study, we aimed to address this gap and disentangle the evolution dynamics of BBCoV-EPI1, a European SARSr-CoV circulating in *Rhinolophus ferrumequinum* bats. The primary hypothesis tested here addressed the influence that a bat’s life history traits would have on BBCoV-EPI1 circulation, on viral evolution, and on lineage emergence. Our results demonstrate that the main drivers of *Sarbecovirus* circulation in *R. ferrumequinum* are ecology, age, and behavior, the latter being strongly shaped by seasonal patterns. After winter torpor, as spring arrives, female bats infected with BBCoV-EPI1 migrate to their reproduction sites. Once there, they clear the virus and its prevalence in the maternity colony steadily declines. Circulation of BBCoV-EPI1 is minimal by early summer, which coincides with the births of all pups. From mid-summer to early autumn, the pups, now juveniles, experience a wave of post-weaning primary infections. Prevalence and modeling allowed for the identification of key epidemiological parameters, with an estimated R₀ of 74 in the colony (MSIRS - see Results). This high value aligns with BBCoV-EPI1’s high transmissibility, which is associated with the prolonged viral shedding (six and a half months) that we observed in the colony. Moreover, the model suggests that immunity to specific BBCoV-EPI1 seasonal variants is short-lived (six weeks), which likely facilitates reinfection and sustained viral transmission within the maternity colony from mid-summer, as females nurse and start weaning their pups. These summer transmissions are mainly driven by primary infections of newly weaned juveniles. Then, during autumn, males and females mate in sites where other species also gather for reproduction, a behavior known as swarming^51^. While adult males, mostly absent from maternity roosts, cannot contribute to infection dynamics in maternity colonies during spring and summer, they gather with females for autumn mating, a period that promotes viral transmission. High prevalence during winter may seem counterintuitive, given the drastic reduction in metabolic activity, body temperature, heart rate, and respiratory rate that occurs during mammalian torpor^52,53^. Such torpor-induced suppression of physiological functions would intuitively be expected to limit viral replication, consistent with previous experiments in controlled cellular environments where low temperatures correlated with reduced viral titers^54–57^. Nevertheless, like other hibernating animals, *R. ferrumequinum* undergoes periodic rewarming during arousal episodes^58^, which may restore physiological functions and temporarily reactivate immune responses, including rapid neutrophils release and increased antibody titers^59^. Moreover, according to the model-derived viral shedding duration of up to 6.5 months, it is possible that individuals infected in October could still excrete the virus in March. Given our results, the winter circulation detected in awake animals may reflect a trade-off between restoration of cellular machinery activity, which promotes viral replication, and recovery of immune functions, which limits viral replication and propagation. While innate immunity may rapidly reactivate after torpor and physiological awakening, we suggest that temperature decrease during torpor might have a negative impact on antibodies’ binding strength and diversity^60^, and thus on adaptive humoral immunity and viral neutralization. Overall, the restricted temporal span (June - October), the structural simplicity of our model and the lack of data do not allow to fully account for biological and behavioral heterogeneity in the colony, such as sex, age, weaning state, and physiological changes associated to the torpor period, susceptible to impact both viral transmission and recovery of individuals. While seasonality and host behavior shape the circulation dynamics of the virus, recombination and single mutations also drive *Sarbecovirus* evolution, leading to heterogeneous evolutionary patterns across genomic regions, a reflection of their functional constraints. Strikingly, we found evidence of positive Darwinian selection in nsp2, nsp3/papain-like protease (PL^pro^) and nsp4, and numerous mutations displayed a notable fitness effect on nsp1-10 of ORF1a, including in nsp5/main protease (M^pro^)/3C-like protease (Fig. 4c (middle), 5c and Supplementary Tables 7 and 12). Proteases are critical for polyprotein cleavage in coronavirus cycles, and several nsps have been described to counteract antiviral immunity in various models. Nsp1, known as a host shutoff factor, selectively inhibits host expression, translation and innate immune functions, thus promoting infectious viral particles production in SARS-CoV, SARS-CoV-2, MERS-CoV and other coronaviruses^61–63^. Native nsp2 has been associated with enhanced viral replication in MHV cell culture, and inhibition of Interferon type I (IFN-I) and Interferon Stimulated Genes (ISGs) responses, as well as increased autophagy, in porcine epidemic diarrhea virus (PEDV)-infected piglets^64,65^. Nsp5/M^pro^ also contributes to inhibit IFN-ß activation by blocking mitochondrial antiviral-signaling protein (MAVS) in SARS-CoV-2^66^. Hence, the strong transient selection of advantageous mutations detected *in vivo* in BBCoV-EPI1 sheds a new light on the implication of nsp1, nsp2 and nsp5/M^pro^ in counteracting antiviral immunity, as it reveals the potential existence of a complex viral - host adaptive balance, for which the mechanisms remain elusive. Besides their proteolytic functions, nsp3/PL^pro^, nsp4 and nsp6 also support viral RNA synthesis and modifications by formation of double-membrane vesicles (DMV), critical for RNA export to the cytoplasm^67,68^. These important functions in polyprotein cleavage, nsps release and viral RNA replication and control, advantageously complement the role of nsp1, potentially through direct interactions in the cytoplasm or on the reticulum membrane. In our model, the positive fitness associated with mutations on these proteins could therefore reflect a better ability to form DMV, enhancing replication and viral load, as well as a more efficient RNA export, thus boosting assembly and egress in response to a changing cellular environment. Taken together, the roles, interactions and temporal dynamics of nsps modifications suggest a constant viral adaptation towards optimal fitness in response to host antiviral defenses that remains to be elucidated. In addition, to decipher the extent and functional impact of protein changes in the evolution of SARSr-CoVs, a central question remains as to the origin of their diversity and the underlying mutations. The error-prone nature of the viral RdRp has long been considered the primary driver of genetic variations, with cell-culture studies showing that proofreading by the nsp14/nsp10 complex strongly constrains polymerase inaccuracy in coronaviruses^69–71^. These models, however, are time-limited and far simpler than natural *in vivo* systems, in which viruses can be followed and sampled repeatedly over years, under realistic selective pressures and against a background of host-driven genetic editing. The evolutionary and cellular processes that shape these mutations - beyond polymerase intrinsic error rates - can therefore be meaningfully assessed only in light of longitudinal studies of viral evolution in natural host populations. The exploration of the BBCoV-EPI1 mutational *spectrum* in bats revealed that host enzymes APOBEC3 and ADAR were accounting for almost 80% of the observed mutations. The notable APOBEC3 editing enzyme, known as an antiviral innate immunity component, and recently described in endemic human coronaviruses (*i.e* NL63, HKU1, OC43, 229E), reflects a long-term virus-host coevolution likely shaped by species-specific enzyme activity^72^. These signatures are weak in SARS-CoV, MERS-CoV, and early SARS-CoV-2 lineages, due to their recent emergence in humans. Therefore, the higher APOBEC3 signature exhibited by more recent SARS-CoV-2 lineages (Omicron - 22B) reflects the ongoing adaptation of this virus to humans. In BBCoV-EPI1, APOBEC3-associated mutation rates are lower than in recent SARS-CoV-2 clades, indicating either a host species specific trait, or a long-term adaptation to *R. ferrumequinum*, or both. Comparison between datasets differing in sampling frequency should be interpreted with caution, as dense longitudinal sampling over extended periods - as proposed here - can capture transient mutations that may subsequently revert, thereby enabling more accurate detection of mutational signatures, particularly those that are deleterious. We found that most of the 55 advantageous mutations are consistent with ADAR activity, and that deleterious mutations were mostly linked to an APOBEC3 profile, suggesting that host editing can have enzyme-specific effects on viral evolution (Fig. 5c). In addition, we found that most S mutations were located in the NTD and associated with variable fitness effects (Supplementary Tables 12 and 13). Though NTD is poorly characterized in the literature, its interaction with the RBD was described in SARS-CoV-2 and seems to exert influence on the S conformation, modulating its receptor-binding capacity, and potentially its tropism^68^. As the NTD can also bind glycans and other cell-surface molecules (*i.e* lectins, kinases)^73,74^, mutations in this domain could affect viral entry efficiency. The T668I deleterious mutation, likely caused by APOBEC3 and located upstream of the S1/S2 cleavage site (Supplementary Tables 12 and 13), may impair recognition by host proteases, thereby controlling viral tropism and fitness. Moreover, several S mutations, such as advantageous Q477H or deleterious P459L that originate from APOBEC3 or ROS, respectively, are located on surface-exposed side chains (Figs. 5c, 6a, 6b and Supplementary Table 13), suggesting they may result from the combined effects of host enzyme activity, oxidative stress, conformation constraints and immune escape pressures. The restoration of infectivity of BBCoV-EPI1 in 2020 is an illustration of this dynamic balance, where beneficial innovations on exposed side-chains (P88S, D246Q, Q477H) are speculated to be associated with immunity escape, compensating for the persistent deleterious mutations (T293I, T668I) located on S2’ and S1/S2 sites with potential conformational and functional changes. In turn, *R. ferrumequinum’s* cellular environment constitutes a bat-specific microecosystem in which ADAR-mediated editing predominates, and APOBEC3-driven innate immune editing and deleterious oxidative stress are markedly weaker than in human cells. Together with strong natural selection, this setting promotes the dynamic evolution of bat sarbecoviruses in nature. The functional interaction between BBCoV-EPI1 RBD and *R. ferrumequinum* ACE2, suggested by AlphaFold 3, was further confirmed by pseudovirus experiments (Figs. 6c, 6e) and aligned with previous works highlighting the critical role of ACE2 in *Sarbecovirus* entry^75,76^. Notably, BBCoV-EPI1 lacks a furin cleavage site at the S1/S2 junction and a TMPRSS2 recognition site, both of which were present in SARS-CoV-2, and instead displays monobasic sites at S1/S2 (TNTR_669_) and S2’ regions (HR_796_). In SARS-CoV-2, furin-mediated cleavage is associated with S conformational changes, leading to RBD exposition and better ACE2 recognition, contributing to its high transmissibility via aerosols^77,78^. The absence of such a site in BBCoV-EPI1 suggests a limited, yet possible, airborne transmission and a favored viral entry via endocytosis. As this entry route is typically associated with enteric tropism, BBCoV-EPI1 may likewise exhibit intestinal tropism, a hypothesis supported by experimental assays on B236, a SARS-CoV-2-related coronavirus, and by another study reporting *Betacoronavirus* titers in bat intestines^79,80^. However, intestinal detection may also reflect swallowed respiratory virus rather than active replication. Thus, robust evidence for active intestinal replication remains limited in the infected bat colonies. Alternatively, the negative fitness effect of T668I, located upstream of S1/S2, could indicate the existence of alternative S1/S2 proteases, which would allow airborne transmission of BBCoV-EPI1. However, the mechanisms underlying its apparent respiratory tropism, suggested by positive detection in oropharyngeal swabs, remain to be clarified^81^. Regarding the potential capacity of spillover to humans, the repulsive ionic charges between hACE2 and BBCoV-EPI1 S (*e.g* Lys480 - His34) make the interaction unlikely so far, which is further confirmed by the inefficient infection of a cell line expressing hACE2 (Fig. 6e). This inability to use hACE2 is widely shared by bat coronaviruses and explains their low isolation rate, as this protein is commonly used in isolation attempts^6,82^. Nevertheless, the dynamics of S mutations observed over a decade of evolution suggest that permissive modifications at these critical amino-acid sites cannot be excluded, albeit they remain difficult to predict. In the sympatric bat species *M. myotis*, ACE2 did not support a BBCoV-EPI1 S-mediated entry into cells. However, modeling experiments still suggest an interaction with BBCoV-EPI1 S. We speculate that this limited affinity is further dampened experimentally, likely because of the absence of species-specific cofactors (proteases, co-receptors, or alternative receptors) in the cellular model. In nature, this may result in transient infections with very limited virus shedding, insufficient for sustained intraspecies circulation. This illustrates that, despite its potential ability to spillover in sympatric bat species, BBCoV-EPI1 still lacks key RBD residues and likely other adaptations needed to emerge and thrive in these new hosts. Considering the conservation of ACE2 among bat species (Supplementary Table 16), this pattern likely extends to other *Myotis* species living in sympatry with *R. ferrumequinum*.

## Methods

### Ethical and legal statements

*Rhinolophus ferrumequinum* is a protected species under European Union and national biodiversity legislation. Sampling was conducted non-invasively, in accordance with relevant conservation regulations and approved permits.

#### SARSr-CoVs dataset compilation, phylogenetic analyses and identity plots

All available human and animal *Sarbecovirus* genomes (n = 216) were downloaded from the NCBI Virus database. Low-quality (>250 Ns), redundant, and highly similar (>99.967% identity) sequences were removed, yielding 21 genomes. These were aligned with MAFFT v7, and the mean pairwise nucleotide sequence identity (MPNSI), relative to the BBCoV-EPI1-P10-G05 reference genome, was computed. *Betacoronavirus* phylogenies were inferred with BEAST 10.4 as previously described^33^. Other phylogenies (accessory/structural proteins, transcription regulatory sequences (TRS), RBM, S1/S2 and S2’cleavage sites), were built with PhyML using SeaView 4 v5.0. Sliding-window identity analysis (w=1,000bp; s=200bp) used BBCoV-EPI1 P10-G05 as the query.

#### Population under study

Guano was collected (n = 2,885) regularly from 2014 to 2024 from a maternity colony of *R. ferrumequinum* bats (Finistère, Bretagne, France). Samples consisted in spatially-separated guano pellets collected beneath roosting sites, each assumed to originate from a single individual. The risk of duplicate sampling was diluted by the large colony size, high fecal output per night, and a guided collection scheme avoiding repeated sampling of the same area. Plus, samples sharing identical consensus genomes were evaluated *post hoc* using plate position and minor variants profiles, to identify potential duplicates. The samples were placed in ammonium sulfate-saturated solution and frozen at -80°C until nucleic-acid extraction was performed as previously described^33^. Ethical approval was not required for this study, as it was based exclusively on non-invasive collection of bat guano and involved no capture, handling, or experimental procedures on live animals, in accordance with French and European regulations.

#### Detection of BBCoV-EPI1

Quantitative real-time RT-PCR targeting the E gene^43^ was performed on 2µL of extract, using SuperScript III Platinum One-Step (InvitrogenTM), on a LightCycler® 480 (Roche). An 8-fold dilution of the Wuhan coronavirus E-gene transcript, obtained from the European Virus Archive, was used as a positive control in each screening batch. Statistical analyses were performed using R studio software. The normality of data distribution was assessed using the Shapiro–Wilk test. Seasonal differences in prevalence were analyzed using the Kruskal–Wallis test, and pairwise Wilcoxon *post-hoc* without adjustment.

#### Epidemiology dynamics

We developed a compartmental maternally-protected-susceptible-infected-recovered-susceptible (MSIRS) model to simulate the transmission of BBCoV-EPI1 in the maternity colony over a 5-month period ranging from July 1st to November 30th. Epidemiological rates were defined using Monolix software (version 2023R1). Details are available in Supplementary Methods 1, and equations are defined in Supplementary Fig. 9.

#### Sequencing

Partial (440 nt) nsp12-RdRp regions of BBCoV-EPI1 were previously amplified by RT-PCR and sequenced for viral characterization^33^. The complete reference genome of BBCoV-EPI1 (P10-G05, 2014) was obtained from a single fecal pellet collected in 2014, and using an untargeted RNAseq approach (https://dx.doi.org/10.17504/protocols.io.14egnr1zzl5d/v1). Based on this reference, we then developed a specific amplicon-based in-house method to deep-sequence viral genomes, consisting in two panels of multiplex primers (Supplementary Table 17), producing 94 overlapping amplicons of an average size of 375 nucleotides covering the whole genome (https://dx.doi.org/10.17504/protocols.io.6qpvr86e2lmk/).

#### Consensus sequences generation

The FASTQ files were QC-filtered using a Phred-score threshold of 20. Cleaned FASTQ files from each sample were mapped to BBCoV-EPI1 P10-G05 reference genome using Bowtie2. The alignments were converted to BAM (Binary Alignment Map) files using Samtools. Only BAM files with a coverage above 85% and nucleotide position with depth above 3X were retained. Variant calling was done with FreeBayes, with standard parameters. The consensus sequences were generated using bcftools and aligned by MAFFT v7.

#### Whole-genome phylogeny

BBCoV-EPI1 whole-genome phylogenetic reconstruction was performed using BEAST v10.5.0. A GTR+I+G substitution model with estimated base frequencies was used, together with an “Uncorrelated relaxed clock” and a constant-size coalescent tree prior. A single MCMC chain was launched for 3x108 iterations to allow convergence and an effective sampling size (ESS) above 200. Recombination was assessed with RDP4. Only breakpoints supported by ≥ 6 algorithms and present in > 2 genomes were used to define the non-recombinant regions. Events with “unknown” parents were excluded if already represented in a known-parent event. Phylogenies of non-recombinant regions were inferred using the whole-genome analyses parameters, with MCMC launched at 1.5×10⁸ iterations. Analyses were performed on full genomes, subgenomic regions, and RDP4-defined regions^83^. Corrected root-to-tip distances (substitution/site) were estimated under GTR+I+G. Sliding-window analyses (w = 1,100 bp; s = 300 bp) used BBCoV-EPI1-P10-G05 and putative recombinant genomes as queries.

### Substitution rates and selective pressure assessment

Variant-calling was performed with LoFreq2* against i) BBCoV-EPI1-P10-G05, to track the majority of mutations over time; ii) the previous year’s genome to compute genomic and subgenomic substitution rates (mutations >50% ÷ region length); iii) each sample’s consensus genome to detect minority variants. dN/dS computation was performed both for subgenomic regions and by sliding window analyses (w: 3,000 bp; s: 300 bp). The log2 fold change (Log2FC) was computed relative to the mean substitution rate of all genomes, for consensus and intra-host diversity. Fitness effect of nonsynonymous mutations was estimated for BBCoV-EPI1 using a pipeline adapted from Bloom and Neher^50^.

#### S and ACE2 modeling

BBCoV-EPI1 S sequences (2014, 2017, 2020) and ACE2 sequences from *H. sapiens*, *M. myotis*, *M. emarginatus*, and *R. ferrumequinum* were retrieved from NCBI. Unstructured regions were removed, and the S ectodomain (residues 1–1141) and ACE2 peptidase domain (residues 20–615) were modeled using AlphaFold3. Models with the highest pTM scores (S: 0.84; ACE2: 0.80–0.90) were analyzed in ChimeraX and structured regions exhibited high pLDDT (local confidence) scores.

#### Cells

A549 cells and A549 expressing ACE2, CD13 or DPP4 (with or without TMPRSS2), were cultured in DMEM with 5% FBS and penicillin (100U/mL) - streptomycin (1100µg/mL) solution (developed *in house*, Synowiec 2021, Lie 2023). Lenti-X 293T cells were cultured in DMEM supplemented with 10% FBS, sodium pyruvate (1mM) and penicillin-streptomycin 100U/mL solution.

#### Plasmids

Codon-optimized ACE2, DPP4, and TMPRSS2 sequences from *R. ferrumequinum* and *M. myotis* were synthesized (GenScript) and cloned into N174 (ACE2, CD13) or pLex307 (DPP4, TMPRSS2) vectors with C-terminal tags (FLAG, His, V5, HA). Human DPP4 lentiviral vector was obtained from Addgene.

#### Establishment of modified cell lines

Lentiviral particles were generated by co-transfecting Lenti-X 293T cells with the packaging plasmid psPAX2, the VSV-G envelope plasmid pMD2.G, and the plasmid encoding the protein of interest. Medium was replaced after 16h, and viral supernatants were collected at 48h and 72h, aliquoted, and either stored at –80 °C or used immediately. A549 cells were transduced with the lentiviral vectors in the presence of polybrene (5 µg/ml) for 16h. After subsequent transduction, cells were maintained in DMEM with selection antibiotics until the non-treated control died (Supplementary Table 18).

#### Assessment of expression

Equalized for total protein concentration cell lysates were separated by SDS-PAGE, transferred to PVDF membranes and probed with specific antibodies. GAPDH was used as load control. The signal was visualized using Immobilon Western Chemiluminescent HRP Substrate. Western Blot analyses were performed using ImageLab software (Supplementary Fig. 10).

#### Pseudotyped virus production and transduction

Pseudoviruses encoding a luminescent reporter, and containing either BBCoV-EPI1 S, VSV-G control, or completely lacking fusion protein, were produced by infecting 293T modified cells with HIV particles (in Lenti-X system; Cells section). Supernatants containing pseudoviruses were collected 48 and 72h after transfection, and used to infect modified A549 cells, in the presence or absence of trypsin (Supplementary Figs 5 and 7). Unbound particles were removed after 16h, and cells were incubated for up to 72 h. Pseudovirus entry was quantified based on luciferase activity. Luminescence was measured using a SpectraMax ID5 microplate reader (Molecular Devices) following Bright-Glo Luciferase Assay (Promega). Data are expressed as normalized Relative Light Units (RLU), calculated as the percentage of the signal obtained with the VSV-G experiments. Statistical analyses were performed using GraphPad Prism 9. The distribution was verified using the Shapiro-Wilk test. According to the results, either one-way ANOVA with Tukey test or Friedman test with Dunn *post-hoc* was used.

#### Confocal microscopy

Fixed cells were permeabilized (15 min, 0.5% Tween-20, Bioshop) and blocked (5% BSA, overnight, 4°C). They were incubated with anti-ACE2, anti-CD13 or anti-TMPRSS2 antibodies coupled with fluorescent secondary antibodies. Nuclear DNA was stained with DAPI. Images were acquired under Zeiss LSM 880 confocal microscope and processed using ImageJ Fiji^84^.

#### Identification of mutational processes

To identify mutational signatures in BBCoV-EPI1 and link them to documented mutational processes, we quantified substitution types in the context of all possible 5’ and 3’ flanking nucleotides. Formatting of plots was performed using the ggplot2 package in R.

#### Data analysis and visualization

All data analyses were performed using R v4.4.2 or bash v3.2.57. Visualizations were produced with ggplot v3.5.1, and identity plots were generated with ggmsa 1.12.0.

## Data availability

Animal and human sarbecoviruses genomes (n = 216) used in the study were downloaded from the NCBI Virus Database. All BBCoV-EPI1 genomes generated in the study have been deposited in the European Nucleotide Archive database under the accession number ERP187933 and embargoed until publication. Accession codes and ID of each sequence are listed in Supplementary Table 19.

## Code availability

In-house scripts generated for this study were written with bash v3.2.57, Python v3.9.5, R v4.4.2 and RStudio v2025.09.1+401, and can be shared upon reasonable request. LightCycler v4.1 (https://www.roche.com) was used for qPCR data collection. Monolix v2023R1 (https://lixoft.com/products/monolix/) was used for epidemiological dynamics modeling. BBTools v39.2 (https://jgi.doe.gov/data-and-tools/bbtools/), Bcftools v1.15 (http://samtools.github.io/bcftools/), Bedtools v2.27 (https://bedtools.readthedocs.io/) and Bowtie v2.5 (http://bowtie-bio.sourceforge.net/bowtie2/index.shtml) were collectively used for sequence data quality control, reads mapping, and generation of consensus sequences. FreeBayes v1.3 (https://github.com/freebayes/freebayes) and LoFreq v2.1.5 (https://csb5.github.io/lofreq/) were used for variant-calling analyses. MAFFT v7 (https://mafft.cbrc.jp/alignment/software/), BEAST v10.5.0 (https://beast.community/), PhyML v3.0 (http://www.atgc-montpellier.fr/phyml/) and RDP4 v101 (http://web.cbio.uct.ac.za/∼darren/rdp.html) were used for phylogenetic and recombination analyses. CLC Main Workbench v26 (https://digitalinsights.qiagen.com/) was used for visualization and annotation of the genome. GraphPad Prism v9.0 (https://www.graphpad.com/) was used for statistical analysis and graphing. ImageJ Fiji and ImageLab v30.1 (https://www.bio-rad.com/) were used for image processing and quantification. Inkscape v1.3.2 (https://inkscape.org/) was used for figure editing and illustration preparation.

## Supporting information

Supplementary_information_concatenated_BBCoV-EPI1_paper

## Acknowledgments

We wish to express our gratitude to Mme Zajec, to pay tribute to the late M. Zajec and to thank the whole family for their support and for their very kind participation in this study. We are grateful to Januel Vouteau and Sophie Levaltier for their contributions to preliminary work, sequencing, and prevalence analyses. We would like to thank Valérie Caro, Mathias Vandenbogaert and Jean-Claude Manuguerra for their precious help at the start of the study. We also thank the Bretagne Vivante Association and the Groupe Mammalogique Breton (GMB) for their support. We also would like to thank Emma Teeling and Albin Fontaine for fruitful discussions, and C.K. for detailed review of the manuscript. Finally, we thank all contributors to the study, wildlife protection associations, field workers, experts, chiropterologists, and volunteers, in particular Matthieu Ménage, Arnaud Le Houédec, Yann Le Bris, Corentin Le Floc’h, the BatLab and the Maglab - Vega team.

## Author contributions

M.A.G conceived the study design. S.L.P., A.V. and M.A.G. acquired the financial support. D.d.R.d.F., S.P., F.T. and M.A.G. carried out sampling. D.d.R.d.F., M.A.G and S.C. performed viral screening. D.d.R.d.F., L.D., M.A.G. and E.L. prepared the NGS libraries and carried out the NGS. D.d.R.d.F., A.F., S.L., B.L. and M.A.G. implemented the bioinformatic analyses or pipelines. D.d.R.d.F. and M.A.G. carried out the phylogenetic and recombination analyses. A.S., A.M. and K.P. developed cell-lines and performed viral entry assays. L.R. and E.D. performed the ACE2-Spike structure modeling and binding studies. B.E. contributed to the development of the MSIRS/SIRS compartment models. D.d.R.d.F. and M.A.G. wrote the manuscript with input from all other authors. All authors reviewed and approved the manuscript.

