## Supplementary_information_concatenated_BBCoV-EPI1_paper for "A 10-year longitudinal study reveals bat-driven SARSr-CoV evolution"

### Supplementary Figures

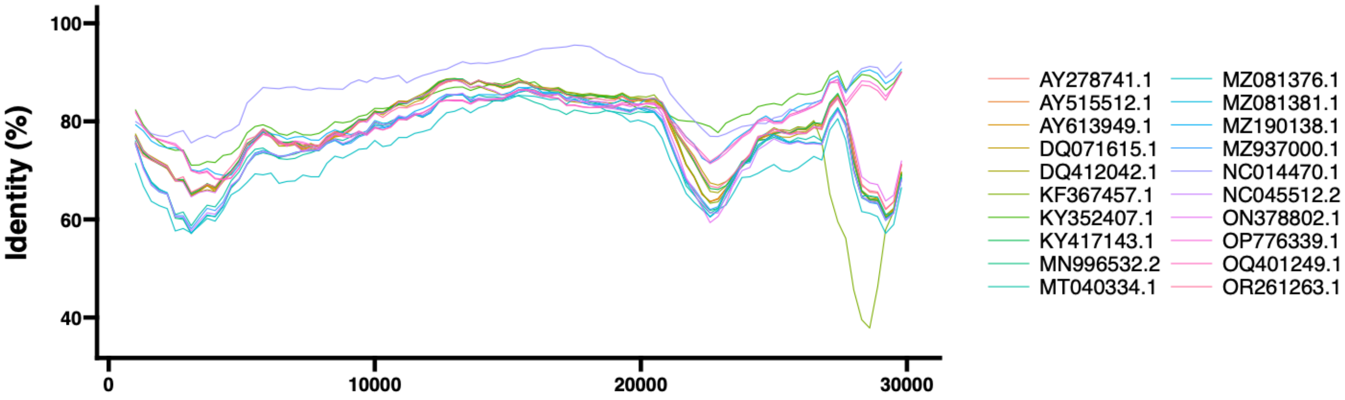


**Figure S1. Identity plot of the 20 human and animal SARSr-CoV genomes mentioned in the study.** BBCoV-EPI1-P10-G05 was used as a query, and each sequence is designated by its NCBI accession number.

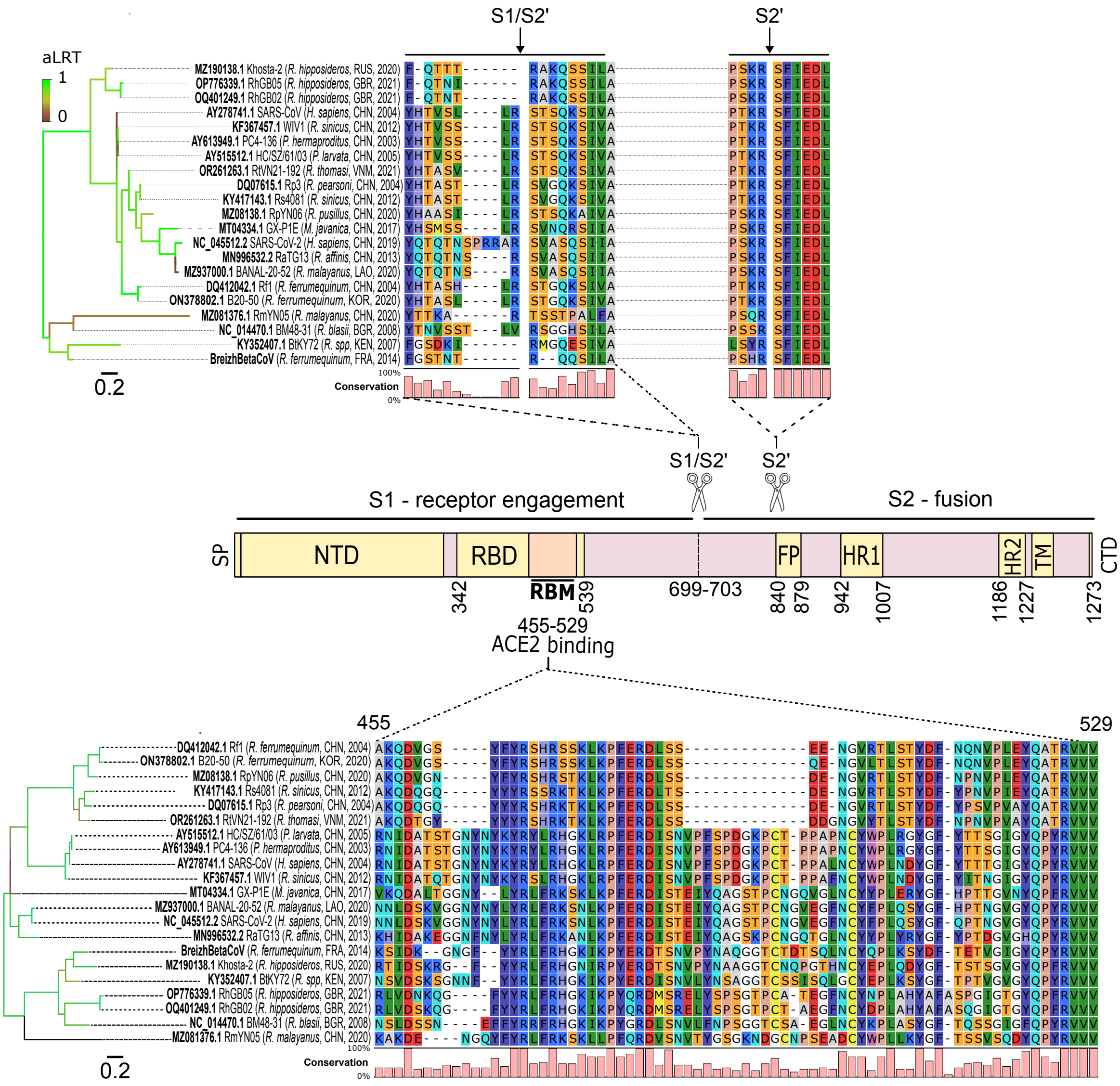


**Figure S2. Amino-acid alignment of S1/S2, S2' and RBM regions of the Spike of the 20 human and animal SARSr-CoV mentioned in the study.** Phylogenies of each alignment were performed with PhyML implemented in SeaView 4. The general time reversible (GTR) model was used, with branches colored according to the likelihood-based measures of branch supports (aLRT). Optimized parameters, such as nucleotide equilibrium frequencies and proportion of invariable sites, were selected. For trees rearrangement and searching operations, both nearest-neighbor interchange (NNI) and subtree pruning and regrafting (SPR) methods were employed.

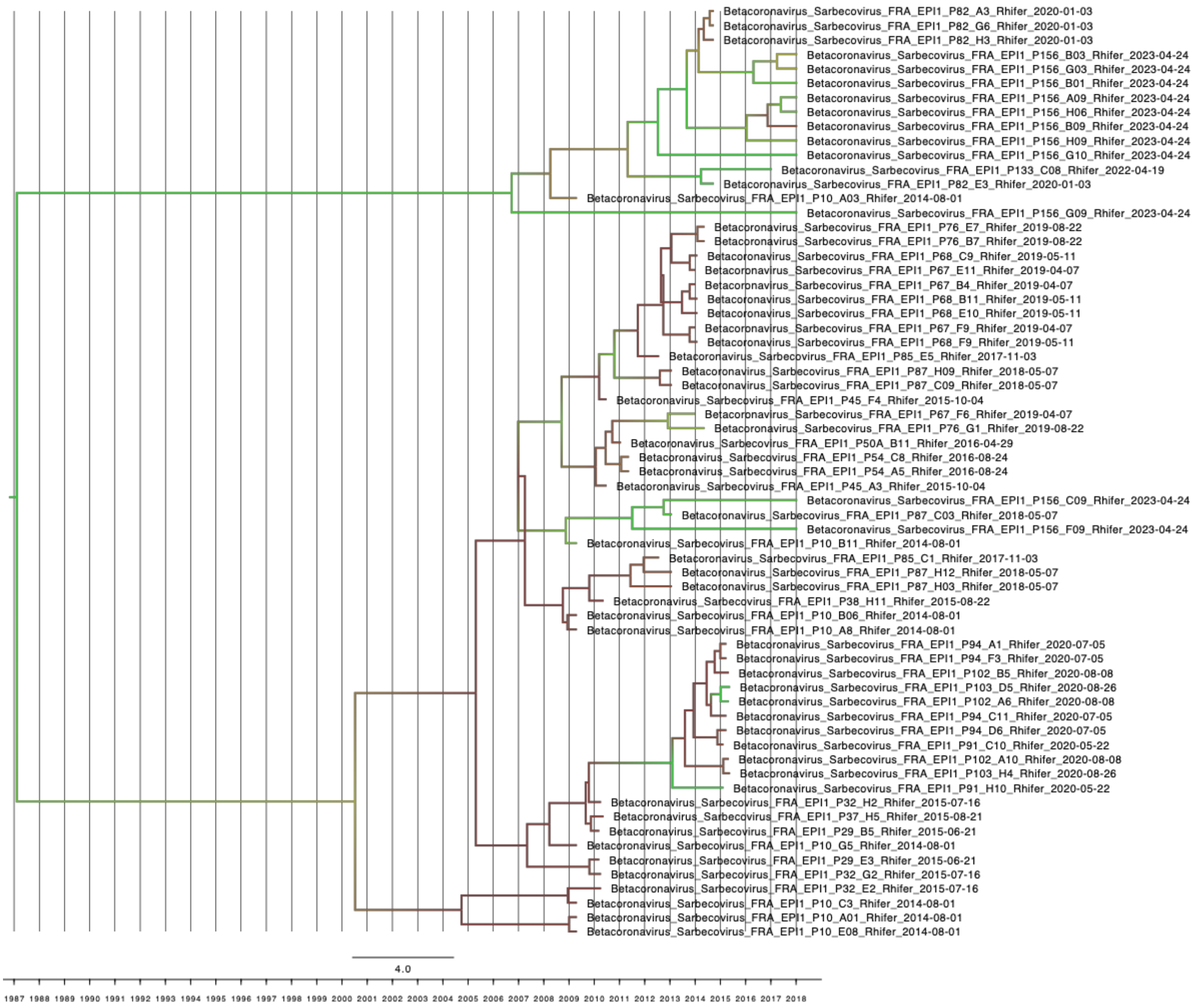

**Figure S3.** Bayesian analysis of region 7 (24,299 – 25,858) of BBCoV-EPI1 genomes. Node support (posterior probability) is shown as a color gradient along the branches.

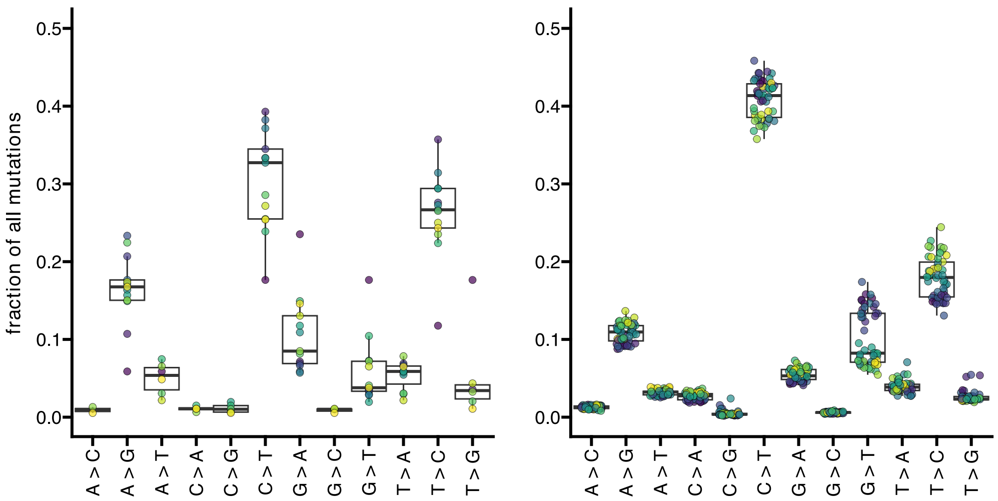


**Figure S4.** Fraction of all mutations of each type for BBCoV-EPI1 (left) and SARS-CoV-2 (right) clades, estimated using four-fold degenerate synonymous sites.

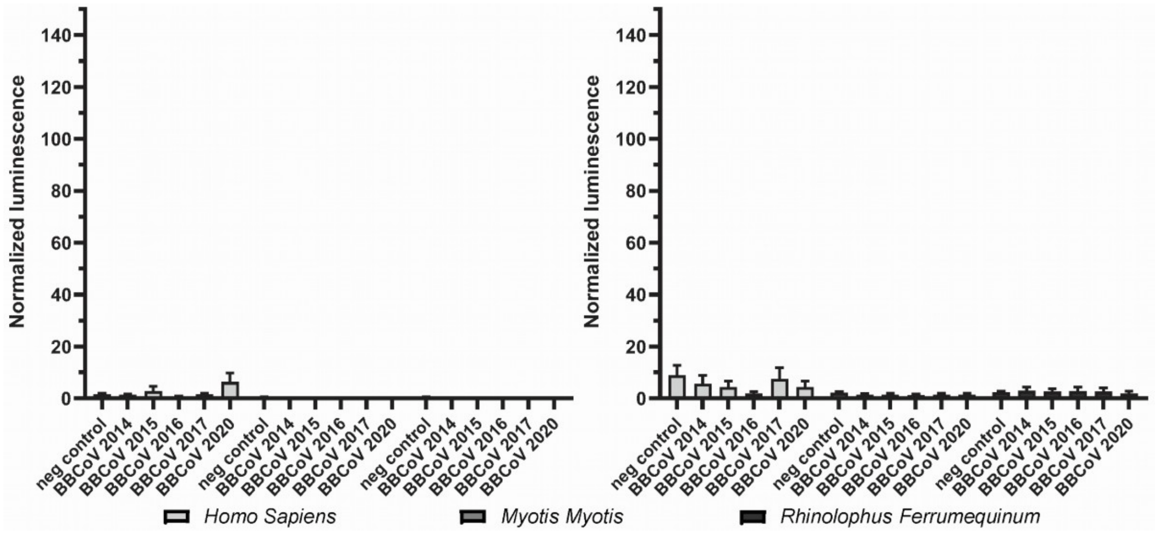


**Figure S5.** Pseudovirus entry to cells overexpressing DPP4, in the presence of trypsin. A549 cells expressing DPP4 alone (left) or both DPP4 and TMPRSS2 (right) were infected with HIV particles pseudotyped with spike proteins based on consensus sequences from the following years, in the presence of trypsin. The data are presented as mean ± SEM from at least three independent experiments performed in triplicate. Three separately prepared sets of pseudoviruses were used.

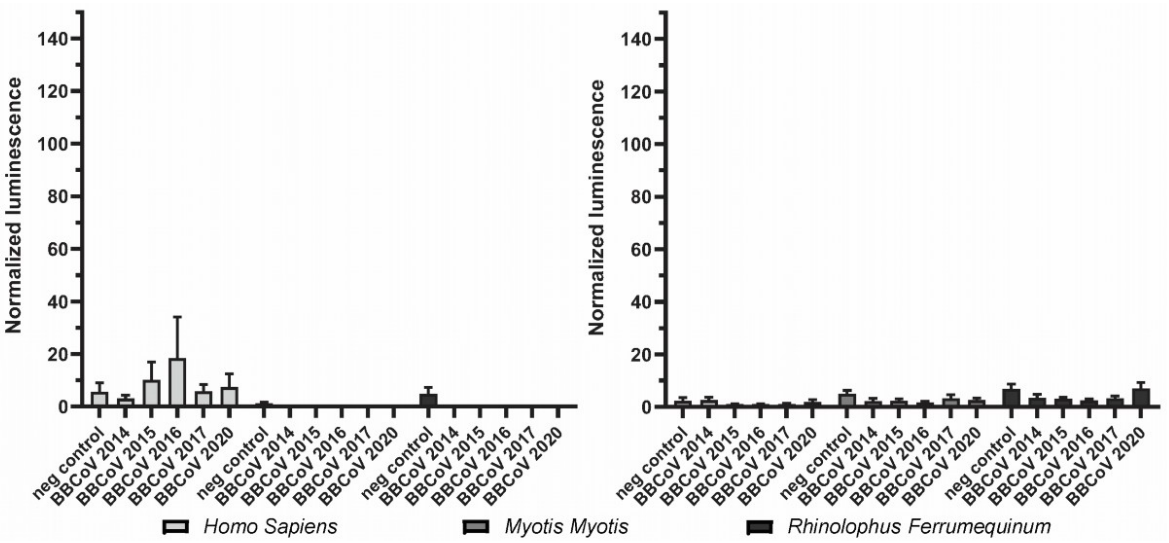


**Figure S6.** Pseudovirus entry to cells overexpressing CD13, in the presence of trypsin. A549 cells expressing CD13 alone (left) or both CD13 and TMPRSS2 (right) were infected with HIV particles pseudotyped with spike proteins based on consensus sequences from the following years, in the presence of trypsin. The data are presented as mean ± SEM from at least four independent experiments performed in triplicate. Three separately prepared sets of pseudoviruses were used.

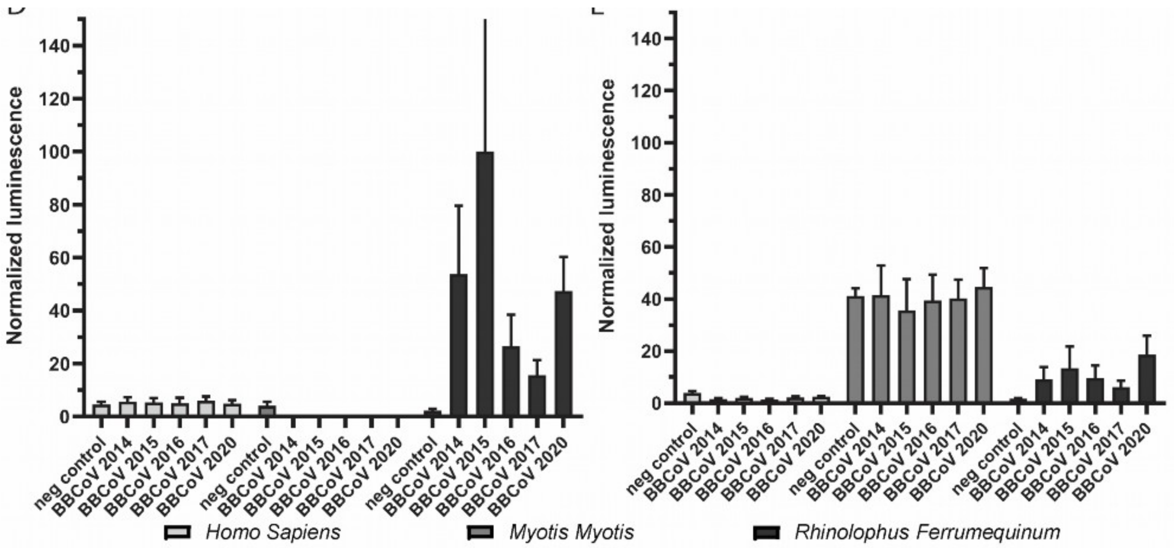


F

| | Neg control | BBCoV 2014 | BBCoV 2015 | BBCoV 2016 | BBCoV 2017 | BBCoV 2020 |
|---|---|---|---|---|---|---|
| Neg control | | *** | *** | ns | ns | *** |
| BBCoV 2014 | *** | | *** | ns | ** | ns |
| BBCoV 2015 | *** | *** | | *** | *** | *** |
| BBCoV 2016 | ns | ns | *** | | ns | ns |
| BBCoV 2017 | ns | ** | *** | ns | | * |
| BBCoV 2020 | *** | ns | *** | ns | * | |

**Figure S7.** Pseudovirus entry to cells overexpressing ACE2, in the presence of trypsin. A549 cells expressing ACE2 alone (left) or both ACE2 and TMPRSS2 (right) were infected with HIV particles pseudotyped with spike proteins based on consensus sequences from the following years, in the presence of trypsin. Two-way Anova with Tukey multiple comparisons test was performed and significance (* p<0.05; ** p<0.01; *** p<0.001) for cells overexpressing *R. ferrumequinum* ACE2 with trypsin are shown. The data are presented as mean ± SEM from at least four independent experiments performed in triplicate. Three separately prepared sets of pseudoviruses were used.

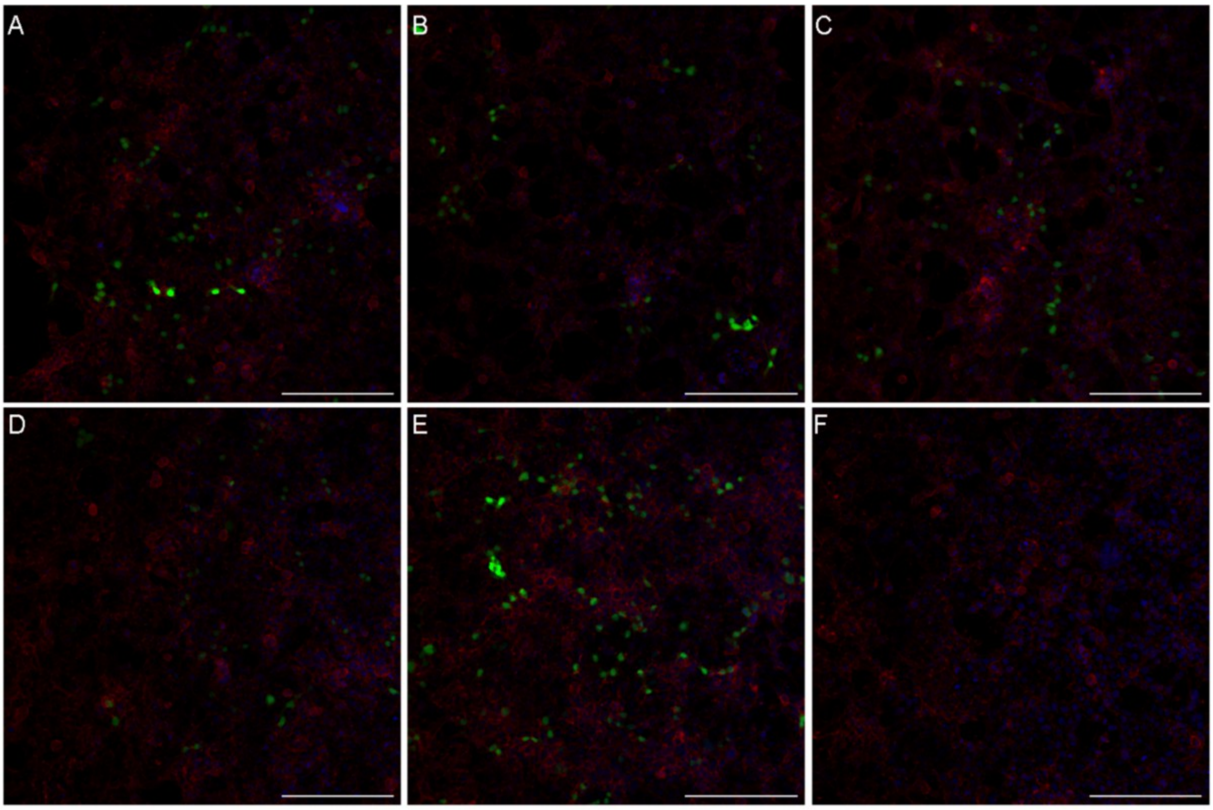


**Figure S8. Pseudovirus entry to cells overexpressing R. ferrumequinum ACE2 & TMPRSS2.** A549 cells expressing ACE2 were infected with HIV particles pseudotyped with spike proteins based on consensus sequences from the following years: A: 2014; B: 2015; C: 2016; D: 2017; E: 2020. F depicts negative control Δenv particles. 72h after infection cells were fixed and in addition to Venus fluorescent protein carried by pseudotyped particles (in green), actin (in red) and DNA (blue) were stained. Cells were visualized under confocal microscope and averaged projection is shown. Bar 200µm.

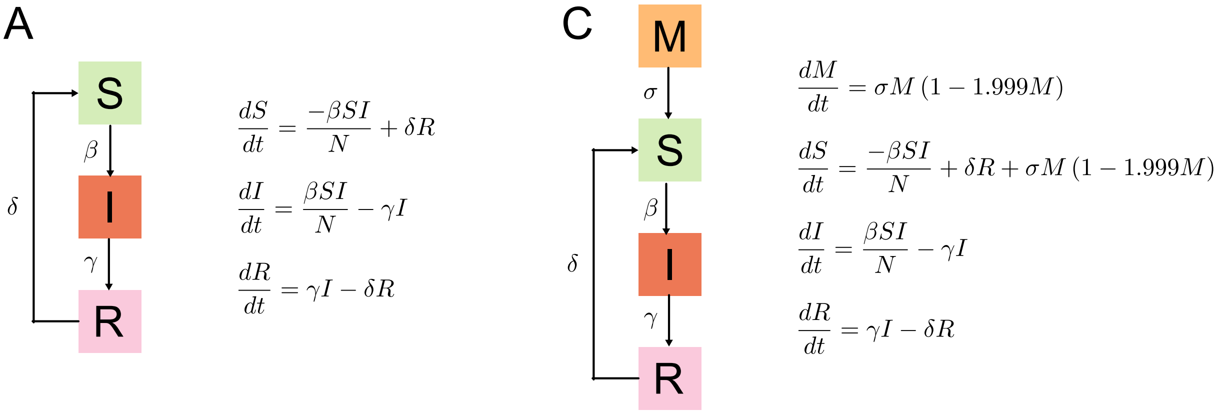


$$\frac{dS}{dt} = \frac{-\beta SI}{N} + \delta R$$

$$\frac{dI}{dt} = \frac{\beta SI}{N} - \gamma I$$

$$\frac{dR}{dt} = \gamma I - \delta R$$

$$\frac{dM}{dt} = \sigma M\,(1 - 1.999M)$$

$$\frac{dS}{dt} = \frac{-\beta SI}{N} + \delta R + \sigma M\,(1 - 1.999M)$$

$$\frac{dI}{dt} = \frac{\beta SI}{N} - \gamma I$$

$$\frac{dR}{dt} = \gamma I - \delta R$$

**Figure S9.** Ordinary differential equations (ODE) of SIRS (left) and MSIRS (right) models.

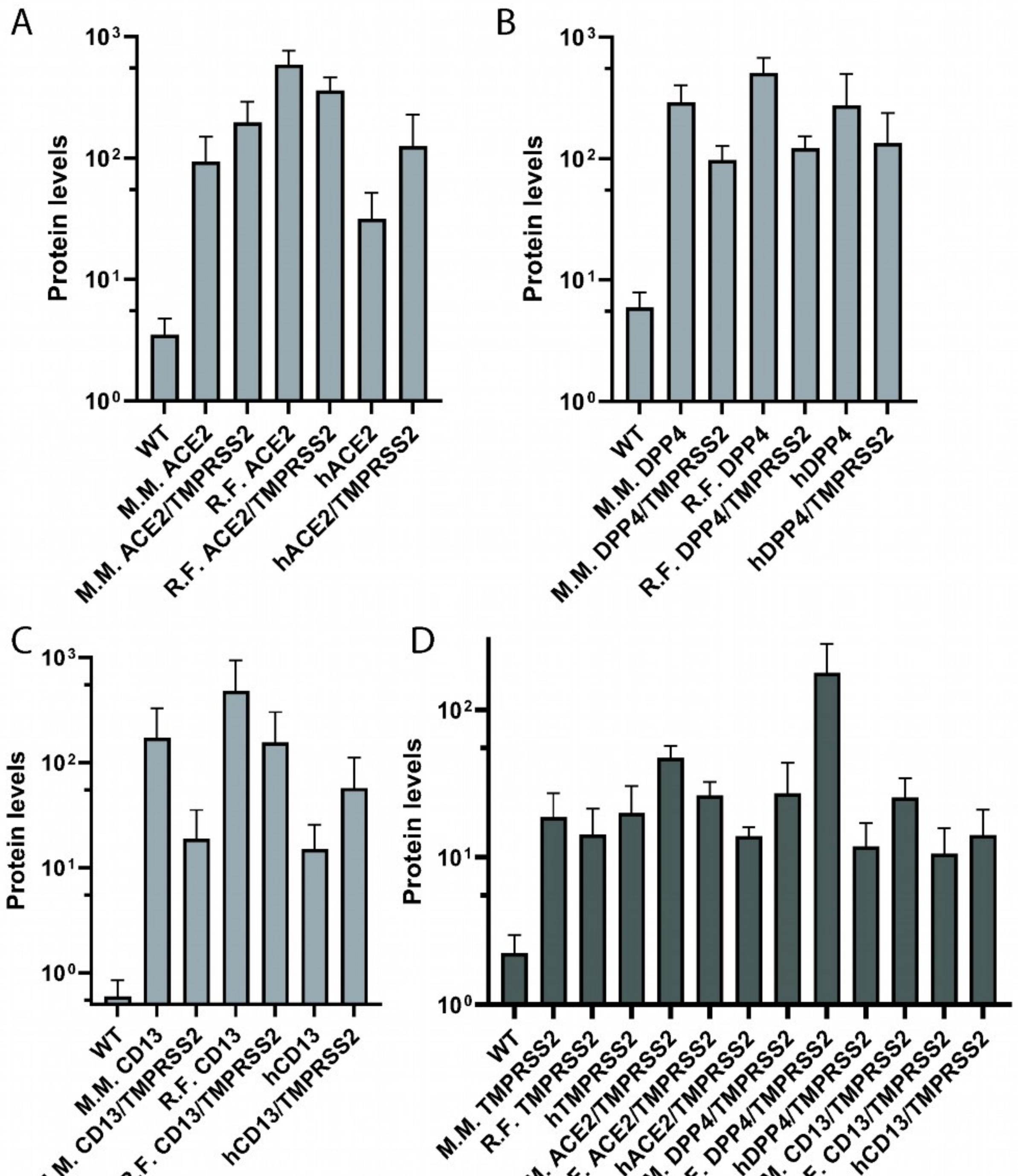


**Figure S10.** Densitometry analyses of modified A549 cells. Levels of ACE2 (A), DPP4 (B), CD13(C) and TMPRSS2 (D) were measured, and the density of bands on collected chemiluminescent blots was calculated using Bio-Rad Image Lab 5 software. The obtained values were normalized to GAPDH levels and are presented as Mean±SEM. Minimum three independently prepared membranes were used. M.M: *Myotis myotis*, R.f.: *Rhinolophus ferrumequinum*, h: *Homo sapiens*.

### Supplementary Tables legends

**Table S1.** MPNSI (%) per genomic region, ordered by MPNSI value. For each region, the closest relative is mentioned.

**Table S2.** Prevalence of BBCoV-EPI1 per sampling plate, sorted by date.

**Table S3.** Variation in BBCoV-EPI1 prevalence and excretion across epidemiological periods. Mean prevalence (%) and mean $\log_{10}$ viral excretion (copies $\mu L^{-1}$) with interquartile ranges (IQR) are shown for each epidemiological period (Low, Intermediate, High). Differences between periods were assessed using Kruskal–Wallis test. Significance levels: ns, not significant; * $p < 0.05$; ** $p < 0.01$; *** $p < 0.001$.

**Table S4.** Estimation of SIRS and MSIRS models parameters using Monolix software. ODE of each model and initial conditions of each compartment are presented in sheets 1 and 3.

**Table S5.** Recombination events detected by RDP4. The columns specify the event number, the number of genomes that exhibit this event, the different algorithms used (R = RDP; G = GENECONV; B = BootScan; M = MaxChi; C = Chimaera; S = SisScan; T = 3Seq), the genomic locations of the breakpoint sites, the putative recombinant associated with the event, and its minor and major parents. Rows in bold indicate the events retained for subsequent analyses.

**Table S6.** Substitution rate (s/s/y) of the 65 BBCoV-EP1 genomes and subgenomic regions sequenced during the study.

**Table S7**. DN/dS *ratio* computed on the major diversity of the 65 BBCoV-EPI1 genomes, from 2014 to 2023, per subgenomic regions.

**Table S8.** Significance of differences in relative rates among the different types of mutations at four-fold degenerate sites identified in SARS-CoV-2.

**Table S9.** Significance of differences in relative rates among the different types of mutations at four-fold degenerate sites identified in BBCoV-EPI1.

**Table S10.** Significance of the differences in relative rates between the different types of mutations at fourfold degenerate sites, between BBCoV-EPI1 and SARS-CoV-2.

**Table S11.** Fraction and relative rates of mutations per clade at 4-four degenerate sites.

**Table S12.** Localization of nonsynonymous mutations identified over the study period, with associated polarity and fitness effect, detected with Bloom & Neher pipeline (Bloom & Neher, 2020).

**Table S13.** Localization of the mutations identified in the BBCoV-EPI1 S over the study period.

**Table S14**. Raw luminescence data from BBCoV-EPI1 S-pseudotyped HIV particle entry assays in A549 cells expressing receptors from different species, with or without trypsin treatment.

**Table S15.** Normalized luminescence data from BBCoV-EPI1 S-pseudotyped HIV particle entry assays in A549 cells expressing receptors from different species, with or without trypsin treatment. Data were normalized to the highest average BBCoV-EPI1 luminescence value (excluding VSV-G control), with BBCoV-EPI1 2015 entry in ACE2-expressing cells from *R. ferrumequinum* set as 100% for graphical representation.

**Table S16.** Alignment of the main residues of SARS-CoV-2, SARS-CoV, PRD-0038 and BBCoV-EPI1-P10-G05 (2014) S, involved in the interaction with the ACE2 receptor. The alignment numbering corresponds to SARS-CoV-2 S, and amino acids are indicated using the one-letter code.

**Table S17.** Sequences of the primers used for the multiplex full-genome amplification of BBCoV-EPI1.

**Table S18.** Supplements needed to maintain populations of introduced proteins.

**Table S19.** Submission of the BBCoV-EPI1 genomes to the European Nucleotide Archive, with accession numbers and IDs of each sequence mentioned.

### Supplementary Methods

**Supplementary Method 1.** Monolix parameters estimation.

To investigate the infection-reinfection dynamics of BBCoV-EPI1 and the contribution of juveniles in its epidemiology, we formulated two mathematical transmission models. In both scenarios, the total population of bats at time t N(t) is considered constant along time (dN/dt = 0).

**SIRS model**

The SIRS model is divided in 3 compartments: Susceptible (S(t)), Infected (I(t)) and Recovered (R(t)). The ordinary differential equations (ODEs) are presented in Supplementary Figure 9.

The proportion of S individuals at the beginning of the scenario was set to 96%, as females are assumed to be infection-permissive at that time, according to both a putative 3-month persistance of protective immunity and field observations showing their waning susceptibility in spring (associated to a decreasing prevalence). The proportions of I and R individuals at the beginning of the scenario were set to 2%, for both. Indeed, prevalence data (Fig. 2a) show that many prevalence estimates in July are ranging from 5% to 10%, with occasional higher values that tend to overestimate the prevalence, resulting in an artifactual LOESS rise and apparent prevalence of about 20% in Fig. 2a.

**MSIRS model**

The MSIRS model consists in the 3 compartments cited above, as well as a Maternally-derived immunised (M(t)) compartment, that aims to take into account the passive immunity of the newborns, induced by transient (or « passive ») protection from maternal antibodies. When maternal antibodies are waned (due to weaning), newborns are included in the S compartment. After being infected, individuals are transferred to the R compartment.

The proportion of M individuals at the beginning of the scenario was set to 50%, as *Rhinolophus ferrumequinum* females usually give birth to a single offspring per year. The proportion of S individuals at the beginning of the scenario was set to 48%. The proportions of I and R individuals at the beginning of the scenario were set to 1%, for both.

We made the hypothesis that BBCoV-EPI1 infection does not induce a lifelong immunity, and that a transfer from R to S compartment is possible. We chose to initiate both simulations from July (when the circulation is the lowest, according to the LOESS regression analysis on sampled data), to November. This timeframe accounts for 63.6% of our sampling effort. Also, it begins at the onset of the maternity period, when *R. ferrumequinum* females give birth, allowing to take into account circulation dynamics from the beginning of this critical stage.

The epidemiological parameters used in the models were defined as follow :

- **the transmission rate β**, represents the proportion of S individuals that are infected by I individuals, per unit of time.
- **the recovery rate γ**, stands for the proportion of infected individuals I that transition to the R compartment, per unit of time. The recovery state involves a complex interaction between specific and innate immunity processes, which are known to be unique in bats.
- **the immunity loss rate δ**, represents the proportion of R individuals that transition to the S compartment, per unit of time.
- **the maternal immunity loss parameter σ** represents the proportion of M individuals, that transition to the S compartment, per unit of time.

***PARAMETERS ESTIMATION USING MONOLIX***

Given our limited knowledge on epidemiological parameters, we used Monolix software (version 2023R1), commonly employed for pharmacokinetic and pharmacodynamic analysis, to estimate epidemiological parameters that best fitted the dynamics observed in our data, in both SIRS and MSIRS scenario.

*Structural model*

As the structural models already present in the Monolix library did not suit our needs, we implemented our own system of ODEs and formatted them in Mlxtran format.

*Initial estimates*

The set typical values for β, γ, δ, and σ were first estimated through manual approximation, in order to align with biological observations of the colony prevalence dynamics. Initial values of β and δ were estimated using the Monolix "Initial estimates" workflow, which employs Bayesian Maximum A Posteriori estimation (MAP), to improve their convergences This approach corresponds to a penalized maximum likelihood estimation based on a prior distribution defined for each parameter. The weight of the *prior* in the estimation is determined by the standard deviation of the prior distribution (see below). γ was fixed to 0.005, to reflect the prolonged viral persistence and extended shedding reported for bat/coronavirus systems, where infections may last substantially longer than in other mammals. In the MSIRS scenario, σ was fixed to 0.1. This choice reflects the progressive transfer of juveniles towards the S compartment, rendering it unfeasible to estimate or to change. Random effects were applied to **γ** in both models, in order to take into account the inter-year variability in disease recovery, which varies according to various factors (*i.e* age, health, sex, immune system maturity). The standard deviation of the random effects (ω) was set to 0.01. The weight of the *prior* in the estimation of the standard deviations of the random effects is linked to the number of degrees of freedom (df) of the student-T distribution. A higher number of df imposes greater constraint on the *prior* in ω estimation. The degrees of freedom were set to 1 for each ω parameter.

*Population parameters estimation*

A lognormal distribution was chosen for each parameter, as it better captures the strictly positive skewness observed in such parameters.

The number of iterations integrated in the burn-in phase (*prior* SAEM) was set to 40, which accounts for at least 10% of the maximum number of iterations in the exploratory phase, set to a minimum of 150 and a maximum of 400. The number of iterations in the smoothing phase was set to a minimum of 100 and a maximum of 200. Convergence of the SAEM algorithm was assessed using Monolix built-in diagnostics, including auto-stop criteria, allowing to automatically detect model convergence, was activated for both phases.

*Standard errors*

The standard errors represent the uncertainty of the estimated population parameters. They are calculated via the estimation of the Fisher Information Matrix, and they can for instance be used to calculate confidence intervals or detect model overparametrization.

The minimum number of iterations of the stochastic approximation algorithm to calculate the Fisher Information Matrix was set to 500, and the maximum was set to 600.

*Likelihood*

A log-likelihood (LL) ratio test was conducted to evaluate the adequacy of the model to the observed data. An importance sampling algorithm was employed to estimate the log-likelihood, as it is better suited for nonlinear mixed effects models compared to the linearization method.

The number of Markov Chain Monte Carlo (MCMC) iterations was set to 1,000,000. Following the likelihood ratio test, -2LL (-2 x log-likelihood), Akaike Information Criterion (AIC), Bayesian Information Criterion (BIC), and corrected BIC (BICc) values were computed and displayed. The AIC of both models were compared in order to assess which model fitted the best the data. The R0 was calculated according to the following formula : $\beta/\gamma$.

**Supplementary Methods 2.** Bioinformatics treatment of sequencing reads.

A. Reads processing and mapping

Raw paired-end Illumina reads were demultiplexed and processed using a custom Nextflow pipeline, available upon reasonable request. Briefly, adapter sequences and low-quality bases ($Q < 20$) were removed using BBduk (BBTools v39.2). Filtered reads were aligned against BBCoV-EPI1-P10-G05 reference genome using Bowtie v2.5. Alignments were sorted into BAM files, and sequencing depth and genome coverage statistics were computed using Samtools v1.15. Genomic positions covered at < 3x were masked with N using Bedtools v2.27, to generate a sample-specific masked reference.

B. Consensus genomes generation

Variants were identified using FreeBayes v1.3, with the following parameters: minimum alternate allele count of 3, minimum alternate allele frequency of 45%, minimum mapping quality of 17, minimum base quality of 20, and variant quality score >20 after filtering with BCFtools v1.15. Consensus genomes were generated by incorporating above-treshold variants into the masked reference using BCFtools v1.15. To characterize intra-host diversity, filtered reads were realigned against their corresponding consensus genomes, and low-frequency variants were identified using LoFreq* v2.1.5. Consensus genome sequences were exported in FASTA format and aligned using MAFFT for downstream phylogenetic analyses.
